# A conserved antigen induces respiratory Th17-mediated serotype-independent protection against pneumococcal superinfection

**DOI:** 10.1101/2023.07.07.548156

**Authors:** Xue Liu, Laurye Van Maele, Laura Matarazzo, Daphnée Soulard, Vinicius Alves Duarte da Silva, Vincent de Bakker, Julien Dénéréaz, Florian P. Bock, Michael Taschner, Stephan Gruber, Victor Nizet, Jean-Claude Sirard, Jan-Willem Veening

## Abstract

Several vaccines targeting bacterial pathogens show reduced efficacy in the context of intercurrent viral infection indicating a new vaccinology approach is required to protect against such superinfections. To find antigens for the human pathogen *Streptococcus pneumoniae* that are effective following influenza infection, we performed CRISPRi-seq in a murine model of superinfection and identified the highly conserved *lafB* gene as virulence factor. We show that LafB is a membrane-associated, intracellular protein that catalyzes the formation of galactosyl-glucosyl-diacylglycerol, a glycolipid we show is important for cell wall homeostasis. Respiratory vaccination with recombinant LafB, in contrast to subcutaneous vaccination, was highly protective against all serotypes in a murine model. In contrast to standard pneumococcal capsule-based conjugate vaccines, protection did not require LafB-specific antibodies but was dependent on airway CD4^+^ T helper 17 cells. Healthy human individuals can elicit LafB-specific immune responses, suggesting its merit as a universal pneumococcal vaccine antigen that remains effective following influenza infection.

**One-Sentence Summary:** Discovery of a universal pneumococcal vaccine protective during superinfection.

## Introduction

*Streptococcus pneumoniae* is a leading cause of bacterial pneumonia and a major cause of death and disability in young children and susceptible adults, including the elderly or immunocompromised. Notoriously, *S. pneumoniae* proves particularly virulent in combination with antecedent influenza A virus infection. Such secondary pneumococcal infections, or superinfections, contribute significantly to excess morbidity and mortality in high-risk groups as highlighted during the influenza pandemics of 1918, 1957, 1968, and 2009^1–4^.

Currently, pneumococcal vaccines are capsule polysaccharide (CPS)-based, such as Prevenar 13^®^, which is composed of 13 CPSs conjugated to a carrier protein together with an aluminum adjuvant and Pneumovax^®^, the pneumococcal polysaccharide vaccine (PPSV) which contains 23 CPSs^5^. Whereas both vaccines elicit CPS-specific antibodies, the conjugated vaccine induces T-cell dependent immunity, which contribute to stronger antibody-mediated protection^6^. While these vaccines are successful in reducing the burden of disease caused by 13-23 serotypes, they do not protect against invasive pneumococcal disease (IPD) caused by non-vaccine serotypes (NVT)^7,8^. There are more than 100 known serotypes of *S. pneumoniae*^9^ and the rapid switching between serotypes, serotype displacement and appearance of non-typeable clinical isolates reduces the efficacy of CPS-based vaccines^10,11^. Importantly, CPS-based vaccines provide poor protection during pneumococcal superinfection following influenza in mice^12,13^. While CPS-based vaccines have shown great protection from IPD caused by serotype-matched pneumococcal strains and likely also contribute to protection following influenza infection, how well they work in this context is unclear from current human vaccine studies^14^. What is clear is that influenza infection contributes to decreased pneumococcal clearance and increased lung injury even in PPSV-vaccinated mice^13^. Conversely, pneumococcal colonization may also impede mucosal immune responses to live attenuated influenza vaccine, including reduced IgA in the nasal cavity and reduced IgG in the human lung^15^.

Thus, there is an urgent need for an efficient vaccine which can cover most virulent pneumococcal strains and provide protection both against primary infection and superinfection. A promising avenue for a universal, serotype-independent vaccine is in the use of immunogenic conserved proteins as protective antigens^5,16–24^. So far, efforts have been focused on surface-exposed pneumococcal proteins as these might be directly recognized by opsonizing antibodies. However, surface-exposed proteins typically show significant strain-to-strain sequence variability because of antigenic variation^25–27^ rendering them prone to vaccine escape and purely protein-based pneumococcal vaccines have still not made it to market. To uncover potential universal antigens, an unbiased genome-wide vaccinology approach is required. Previous attempts have used transposon insertion sequencing (Tn-seq) to identify pneumococcal antigens^28,29^. While successful, these approaches identified non-essential genes encoding for variable surface-exposed proteins that suffer from the limitations outlined above. Here, employing CRISPR interference (CRISPRi) that allows the interrogation of essential genes^30^, we searched specifically for conserved genes highly important for bacterial survival during superinfection. We show that one of our hits, LafB, a highly conserved membrane-associated protein, is an essential virulence factor. Importantly, recombinant LafB provides broad Th17-specific protective immunity paving the way for a universal, capsule-independent, pneumococcal vaccine.

## Results

### CRISPRi-seq identifies LafB as novel pneumococcal virulence factor for influenza superinfection

We previously built a doxycycline-inducible genome-wide CRISPRi library that targets 99% of genetic elements present in the virulent serotype 2 D39V *S. pneumoniae* strain^31^. By sequencing and quantifying sgRNAs in doxycycline-free or -supplemented conditions (to induce dCas9), the relative fitness of each targeted feature can be determined^30^. Using this CRISPRi-seq approach in mice fed doxycycline-containing food, we confirmed pneumococcal capsule as an important virulence factor during superinfection^31^. To more precisely control *in vivo* dCas9 expression, doxycycline levels in serum and epithelial lining fluids (*i*.*e*. bronchoalveolar lavage) were optimized following intraperitoneal (i.p.) injection in mice. A novel *ex vivo* CRISPRi-based luciferase assay found as little as 4 ng/ml doxycycline repressed luciferase transcription >15-fold (**Figure S1**); i.p. injection of 5 mg/kg of doxycycline adequately activated the pneumococcal CRISPRi system in the lung.

Next, mice were infected intranasally (i.n.) with H3N2 influenza virus followed at day 7 by i.n. infection with the *S. pneumoniae* CRISPRi library. dCas9 was induced by doxycycline and compared to mock (vehicle) control (**Figure 1a**). The CRISPRi-seq screen confirmed the capsule operon as critical for pneumococcal survival in the host; in contrast, the *in vitro* essential gene *metK* was dispensable *in vivo*^31^(**Figure 1b**). To pinpoint conserved *S. pneumoniae* genes with important virulence functions in the context of influenza superinfection that could be promising vaccine candidates, we plotted the fitness values of each clone across *in vitro* and *in vivo* conditions. This analysis identified sgRNA0370 targeting the gene *spv*_*0960* (*lafB*), previously unnoticed in Tn-seq experiments, to be significantly underrepresented *in vivo* following doxycycline induction (**Figure 1b, Supplementary Table 1**).

**Figure 1.**
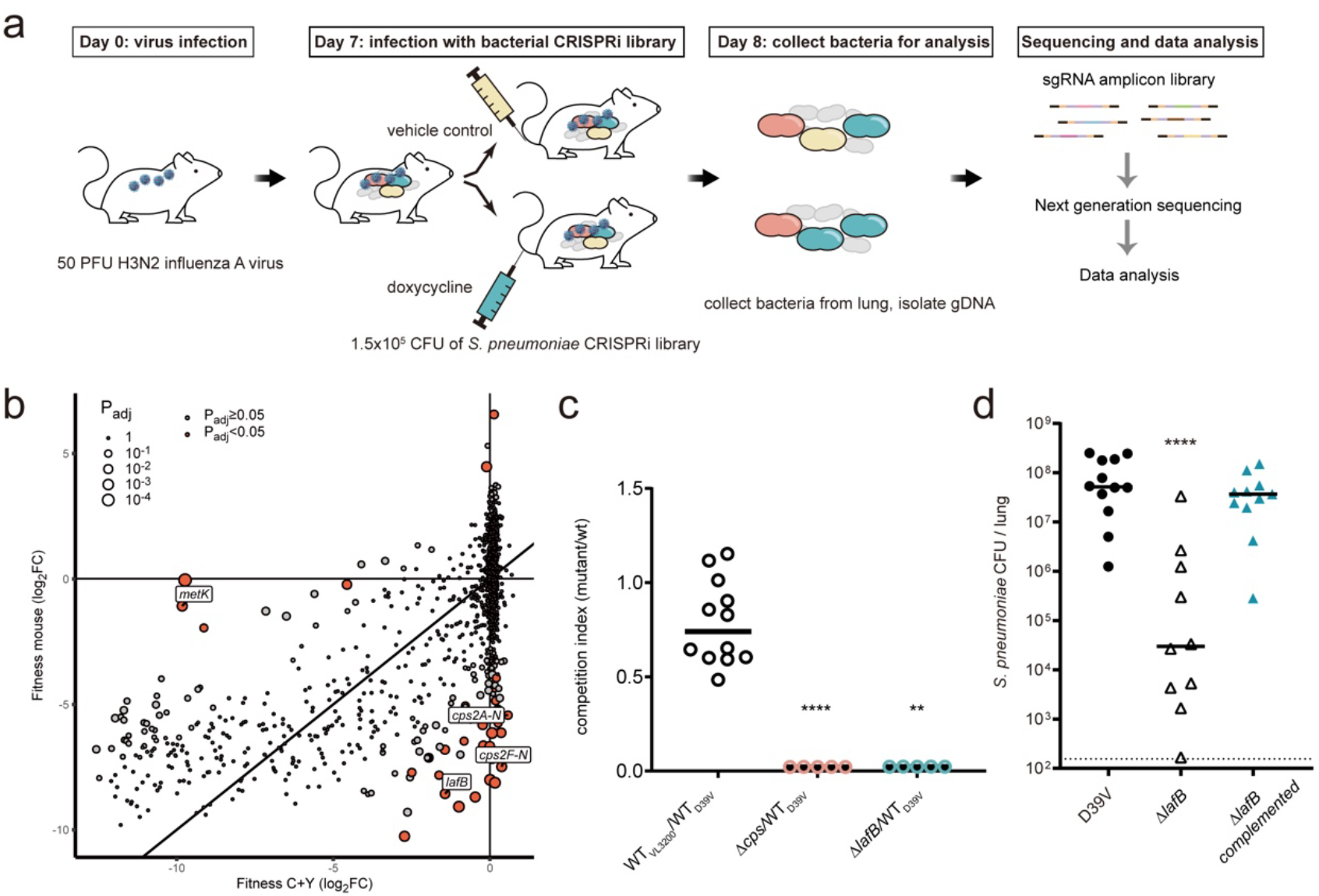
LafB is an essential virulence determinant in pneumococcal pneumonia following influenza infection. (**a**) Workflow of the CRISPRi-seq screen using injected doxycycline. Mice were inoculated intranasally with the genome-wide pneumococcal CRISPRi library. (**b**) CRISPRi-knockdown of the capsule operon (*cps2A-N, cps2F-N*) and *lafB* results in reduced fitness *in vivo* (mouse) compared to *in vitro* (C+Y medium). (**c**) Competition index of individual mutants, compared to wild type (WT) D39V. The Δ*lafB* and Δ*cps* mutants were outcompeted by the WT strain. Strain VL3200 is similar to WT but contains an erythromycin resistance marker at a neutral locus to allow for selection. Each data point represents the lung CFU count at day 8 of a single mouse infected with flu at day 0, and a ratio 1:1 of mutant and WT strain at day 7. (**d**) Validation study of sgRNA0370 target, *lafB*. The Δ*lafB* mutant was attenuated in establishing lung infection. Ectopic expression of *lafB* complemented the phenotype. Kruskal-Wallis testing was used to compare groups.

To validate the CRISPRi-seq screen, *lafB*-deleted and complemented mutants were constructed (**Figure S2a-b**). Competition assays were conducted 7 days post influenza infection with wild type *S. pneumoniae* paired with a *lafB* mutant or a *cps* mutant (avirulent control) in a 1:1 ratio. *S. pneumoniae* lacking LafB were outcompeted by wild type bacteria suggesting a major role of *lafB* for replication in the host (**Figure 1c**). These results were confirmed in single strain superinfection experiments, as *lafB* mutant bacteria had significantly reduced in lung bacterial counts compared to the wild type or *lafB*-complemented strains (**Figure 1d**). Invasive disease, assessed by splenic dissemination, was likewise attenuated in animals infected with the *lafB* mutant (**Figure S2c-d**), indicating LafB is essential for pneumococcal virulence.

### LafB is an intracellular membrane-associated protein involved in cell wall homeostasis

Lipoteichoic acid anchor formation protein B (LafB, 347 amino acids, 40 kDa)^32^ is highly conserved among pneumococci (>96% amino acid identity in all sequenced pneumococci) and closely related members of the Streptococcus *mitis* group (**Figure S3a-b**) and has been implicated in the production of galactosyl-glucosyl-diacylglycerol, a glycolipid of unknown function (**Figure 2a**)^33,34^. Incubation of recombinant LafB with α-monoglucosyldiacylglycerol (mGlc-DAG) and UDP-Galactose followed by mass spectrometry, demonstrated the production of UDP (**Figure S4**), establishing that LafB is a diglucosyl diacylglycerol synthase, as proposed previously^35^. Additionally, *lafB*-deficient pneumococci have a slight reduced susceptibility to penicillins^33^, but increased susceptibility to daptomycin and acidic stress^34,36^. Prior western blot analysis found LafB co-purifies with the membrane fraction^33^, However, while our structure prediction using RoseTTAFold^37^ demonstrates the Rossmann-like domain of GT-B glycosyltransferases^38^, no clear transmembrane domains were detected (**Figure 2b)**. Overlay of our predicted model of LafB to a crystalline structure of a structurally related GT-B glycosyltransferase, *Mycobacterium tuberculosis* PimA (Pdb 2GEK)^39^, showed good agreement, albeit with deviations in the active site cleft, bespeaking the different substrate specificities of the two proteins (**Figure S3f**).

**Figure 2.**
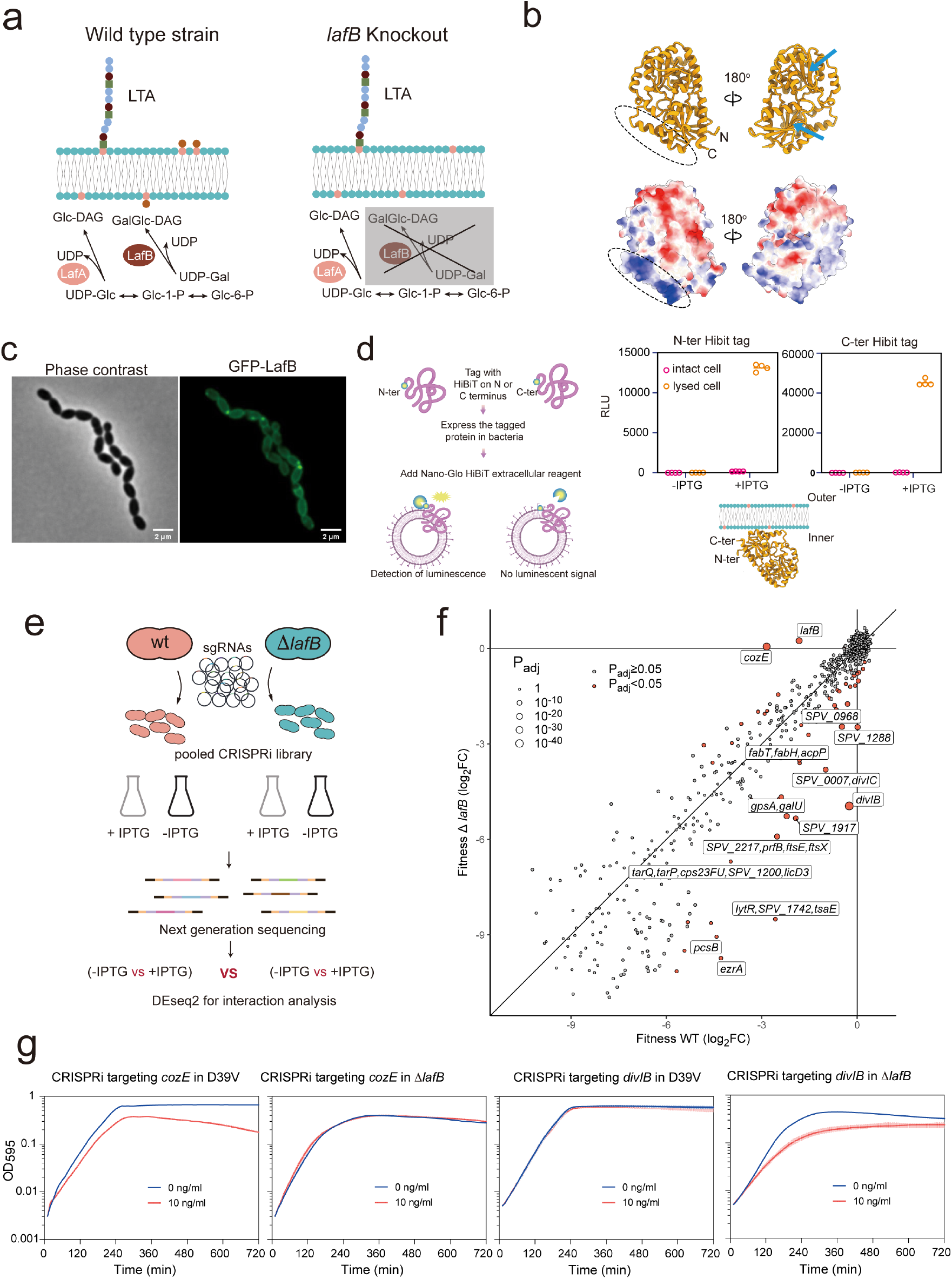
LafB is a membrane-associated galactosyl-glucosyl-diacylglycerol synthase with a pleiotropic role in cell division and cell wall homeostasis. (**a**) LafB is a glycosyltransferase encoded in the same operon with another glycosyltransferase, LafA. LafA catalyzes the synthesis of glucosyl-diacylglycerol (Glc-DAG), which provides the anchor for lipoteichoic acids (LTA). LafB catalyzes the addition of galactose onto Glc-DAG synthesizing GalGlc-DAG (**Fig. S4**). In the absence of LafB, no detectable of GalGlc-DAG can be found in *S. pneumoniae*. (**b**) The predicted structure of LafB by RoseTTAFold. Negative and positive electrostatic potentials are colored red and blue, respectively. The two blue arrows point to the active units. No transmembrane domain was identified. (**c**) GFP was fused to the N-terminus of LafB (GFP-LafB) and fluorescence microscopy analysis showed a membrane localization. (**d**) HiBiT assays showed that both N- and C-termini are localized inside the cytoplasm. LafB was tagged with the HiBit tag at its N- or C-terminus and placed under an IPTG-inducible promoter. Only when the HiBiT is present outside the cell, it can interact with the complementary LgBiT and reconstitute the luminescent NanoBiT enzyme (see Methods). Luminescence (relative light units; RLU) is recorded with a microplate reader. (**e**) The workflow of CRISPRi-seq in wildtype (WT) D39V and *lafB* knockout mutant (Δ*lafB*) to identify the gene interaction network. (**f**) Comparison of fitness cost of gene depletion by CRISPRi between wild type and Δ*lafB* mutant. The sgRNAs showing significant fitness cost between WT and Δ*lafB* are colored in orange and their targets are labelled. (**g**) Growth curve of WT and Δ*lafB* mutant with doxycycline inducible-CRISPRi targeting *cozE* and *divIB* confirmed the positive interaction of LafB/CozE and negative interaction of LafB/DivIB. Strains were pre-cultured to mid-exponential phase, diluted 1:100 in C+Y medium with (10 ng/ml) or without (0 ng/ml) doxycycline. Turbidity of the cell culture is monitored by a microplate reader at 595 nm (OD595) every 10 min. Average of 3 replicates is presented. Shadow showed the range of the measured OD595.

To pinpoint LafB cellular localization, we constructed a functional LafB-GFP fusion expressed from its native locus (**Fig. S3c-e**) and performed fluorescence microscopy on live *S. pneumoniae*. As shown in **Figure 2c**, LafB-GFP demonstrates clear membrane-associated localization. Split complementation luciferase assays for topology showed that both LafB termini reside in the cytoplasm (**Figure 2d**). These data support our structural model of LafB as an intracellular protein that is membrane-associated via hydrophobic and charge interactions with the cytoplasmic membrane.

To gain additional insight into *lafB* mutant virulence attenuation, we performed a genome-wide synthetic lethal screen by introducing a sgRNA library containing 1499 unique sgRNAs into the Δ*lafB* mutant background, then grew bacteria under laboratory conditions where *lafB* is not essential (**Figure 2e**). As shown in **Figure 2f**, the gene encoding the division protein DivIB^40^ becomes more essential in a *lafB* mutant background, suggesting that galactosyl-glucosyl-diacylglycerol plays a role for efficient cell division. The gene *cozE* (aka *cozEa*) encoding a known regulator of penicillin-binding protein Pbp1A^41^ becomes less essential in absence of *lafB* (**Figure 2f**). This genetic interaction may reflect prior findings that *cozE* mutants have deranged Pbp1A activity causing cell lysis^41^. Since *lafB* mutants have reduced Pbp1A levels^33^, a double *lafB*/*cozE* knockdown alleviates the *cozE* single mutant phenotype. Testing individual knockdowns of *divIB* and *cozE* validated the screen (**Figure 2g**). These pleiotropic effects of *lafB* deletion on membrane and cell wall physiology likely underpin the attenuation of virulence of the Δ*lafB* mutant (**Figure 1d**).

### Vaccination with LafB induces antigen-specific adaptive immune responses

To establish whether LafB is a protective vaccine antigen, we cloned *S. pneumoniae* D39V *lafB* and produced the protein in *E. coli* (**Figure S4a**). Recombinant LafB was formulated with alum as adjuvant for subcutaneous (s.c.) immunization, or with the recombinant *Salmonella enterica* serovar Typhimurium flagellin FliC_Δ174-400_ as a mucosal adjuvant^42–44^ for i.n. immunization. Adaptive immune responses specific for LafB were tested in mice on day 28 after a prime-boost vaccination (**Figure 3a**). A strong LafB-specific antibody response (IgG, IgM but no IgA) was observed for s.c. vaccinated animals in serum and broncho-alveolar lavages (BAL), respectively (**Figure 3b and Figure S5a-d**). In contrast, LafB-specific antibodies were weakly elicited in mice vaccinated via the i.n. route. When immune cells from lung, spleen and mediastinal lymph nodes (MdLN) were stimulated *ex vivo* with LafB antigen, cytokines associated with Th1 (IFN*γ*), Th2 (IL-13), and Th17 (IL-17/IL-22) were produced in response regardless the vaccination route (**Figure 3c** and **Figure S5e-f)**.

**Figure 3.**
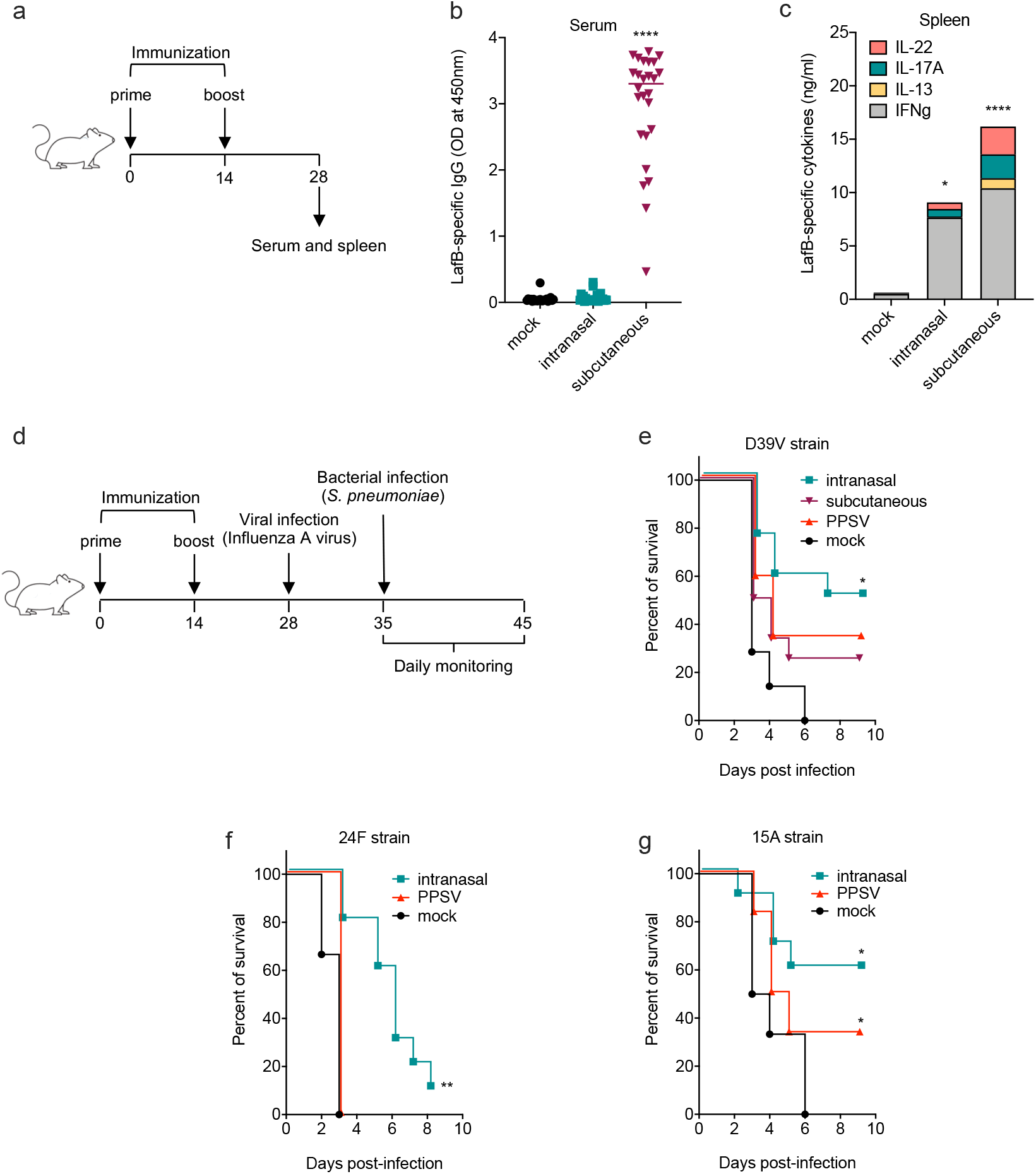
Intranasal vaccination with LafB protects mice against pneumococcal disease in a serotype-independent manner. C57BL/6 mice (n=6-26) were immunized at days 0 and 14 with LafB via intranasal (flagellin-adjuvanted) or subcutaneous (alum-adjuvanted) route, a commercial PPSV vaccine, or left untreated (mock). (**a**) Vaccination and immune response timeline. (**b-c**) Immune responses at day 28. (**b**) LafB-specific antibody response. Sera were collected and levels of LafB-specific IgG were determined by ELISA. Plots represent values for individual mice as well as median. (**c**) LafB-specific T cell response. Spleen cells were stimulated 72 h with LafB and cytokine levels in supernatant were determined by ELISA. Results are expressed as median. Statistical significance (*P<0.05, **** p<0.0001) was assessed by one-way ANOVA Kruskal-Wallis test with Dunn’s correction compared to the mock group. (**d-g**) Analysis of vaccine protective efficacy. (**d**) Vaccination and challenge timeline. Vaccinated mice were infected with H3N2 influenza A virus at day 28 and were challenged at day 35 intranasally with *S. pneumoniae* strain serotype 2 D39V (**e**, 5×10^4^ CFU), serotype 24F (**f**, 10^3^ CFU), or serotype 15A strain (**g**, 5×10^4^ CFU). N= (**e-g**) Protection was assessed by monitoring survival. Statistical significance (*P<0.05, **** p<0.0001) was assessed by Mantel-Cox test compared to the mock group. Each group had at least 10 mice.

### Intranasal vaccination offers broad protection across serotypes

Vaccinated animals, including PPSV-immunized animals, were infected on day 28 with H3N2 influenza virus and superinfected on day 35 with the *S. pneumoniae* serotype 2 strain D39V (**Figure 3d**). All mice receiving mock immunization succumbed to disease after infection. Forty percent of mice vaccinated with PPSV, which includes the CPS from serotype 2, were protected against pneumococcal challenge (**Figure 3e**). Mice vaccinated via the i.n. route with flagellin-adjuvanted LafB outperformed both subcutaneous- and PPSV-vaccinated animals, with 60% mouse survival. LafB standalone i.n. vaccination was poorly effective (**Figure S5g-h**). Mice immunized i.n. with the flagellin-adjuvanted irrelevant antigen ovalbumin (OVA) were not protected (**Figure S5i-j**). These data demonstrate that LafB is a protective antigen against pneumococcal infection when formulated with an intranasal adjuvant.

Western blotting showed that serum of animals vaccinated with LafB from serotype 2 strain D39V recognized all tested strains representing serotypes 1, 3, 4, 5, 9V, 11A, 15A, 19F, 23A, 23F, 24F and 35B, corroborating the high conservation of the LafB protein sequence across pneumococci (**Figure S4e**). Since the introduction of the CPS-based vaccines, NVT are becoming prevalent^7,11^, in particular serotypes 15A and 24F^7,45,46^, which are not included in PPSV (that does however contain 15B, which is poorly cross-reactive to 15A)^47^. As shown in **Figure 3f-g** and **Figure S6**, flagellin-adjuvanted LafB vaccination significantly protected mice against infection with NVT 15A and 24F, in stark contrast to mice vaccinated with PPSV, which only offered slight protection against serotype 15A. In contrast to PPSV controls, LafB-vaccinated mice completely cleared pneumococcal bacteria (**Figure S6e-f**), supporting a role for LafB as a universal vaccine antigen to confer sterilizing protection against pneumococcal infections.

### Protection against pneumococcal superinfection is mediated by Th17 immunity

Th17 CD4^+^ T lymphocytes that are functionally characterized by the expression of the retinoid orphan receptor g t (RORgt) and the production of IL-17A, are essential for mucosal protection against pneumococcal nasopharyngeal colonization and infection^48–51^. To get more insight on the mechanisms of vaccine protection, mice were immunized i.n. with flagellin-adjuvanted LafB, and then infected with influenza virus. Immunoprotective responses were monitored starting from day 35, a time when viral infection impairs the innate and cell-mediated immune responses^52–56^ (**Figure 4a**). In this context, cells isolated from spleen, MdLN or lung from the i.n. vaccinated animals secreted IL-17A after *ex vivo* stimulation with LafB antigen (**Figure 4b**), indicating that influenza infection did not disturb the capacity of the vaccine to stimulate IL-17A. Moreover, vaccination did not intrinsically alter viral replication nor the virus-induced pro-inflammatory response when compared to mock or s.c. immunization, as measured by the viral RNA copy number and major markers for lung inflammation (**Figure S7**). In contrast to wild type animals, *Il17a*-deficient mice were not protected against superinfection by *S. pneumoniae* after the i.n. vaccination (**Figure 4c**). Together, these data show that IL-17A is a major effector cytokine of immunoprotective response induced by LafB i.n. vaccination.

**Figure 4.**
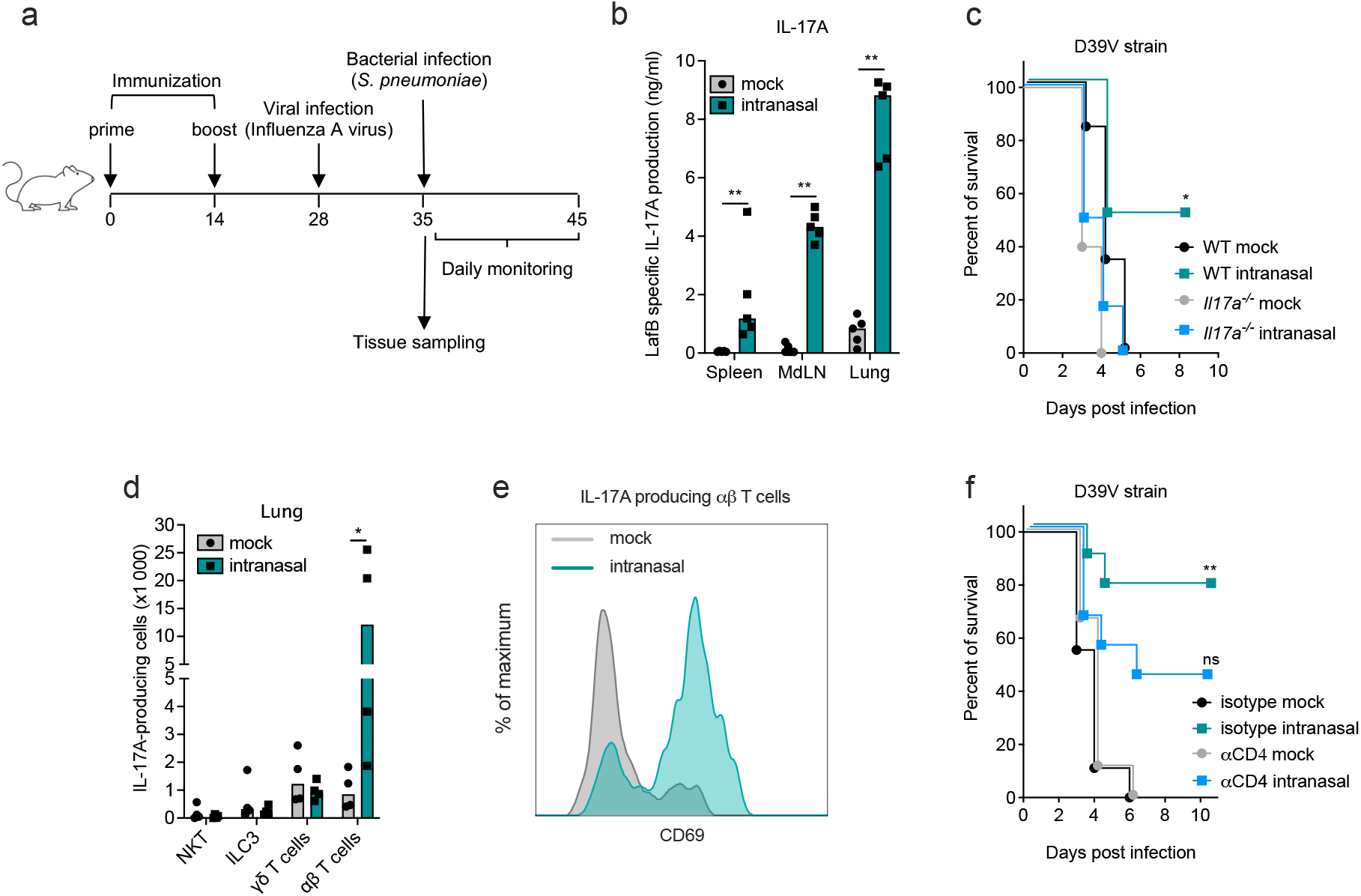
Protection is mediated by Th17 lymphocytes with TRM features. C57BL/6, *Rorc(t)-Gfp*^*TG*^ or *Il17a*^*-/-*^ mice (n=4-10) were immunized at days 0 and 14 with flagellin-adjuvanted LafB by intranasal route or left unvaccinated (mock) and infected with H3N2 influenza A virus at day 28. At day 35, the immune responses of virus-infected animals (**b, d, e**) were analyzed, or the animals were superinfected with *S. pneumoniae* strain D39V (5×10^4^ CFU) to monitor survival (**c, f**). (**b**) LafB-specific IL-17A secretion. Spleen, MdLN and lung cells from C57BL/6 animals were collected and stimulated 72h with LafB antigen. IL-17A levels in supernatant were determined by ELISA. Plots represent values for individual mice as well as median. Statistical significance (**P<0.01) was assessed by Mann-Whitney test compared to the mock group. (**c**) Vaccine protection is abolished in *Il17a*^*-/-*^ mice. Statistical significance (*P<0.05) was assessed by Mantel-Cox test compared to the mock group. (**d-e**) RORgt- and IL-17A-producing lung cells in *Rorc(t)-Gfp*^*TG*^animals. (**d**) Analysis of Natural Killer T (NKT) cells, group 3 innate lymphoid cells (ILC3), TCRγδ T cells, and conventional αβ T lymphocytes. Plots represent values for individual mice as well as median. Statistical significance (*P<0.01) was assessed by Mann-Whitney test compared to the mock group. (**e**) Expression of CD69 marker on lung CD4^+^ Th17 cells. (**f**) Protection requires CD4+ T cells. To this end, mice were treated intraperitoneally at day 34 with CD4-specific depleting antibodies or control isotype, infected at day 35 with D39V and protection was assessed by monitoring survival. Statistical significance (** p<0.01) was assessed by Mantel-Cox test compared to the mock group.

Focusing on IL-17A-producing cells in the lungs (**Figure 4d-e**), we found the main cells producing ROR γt and IL-17A after i.n. vaccination and influenza virus infection were conventional CD4^+^ T lymphocytes expressing TCRαβ, *i*.*e*. Th17 lymphocytes. Other innate lymphocytes, such as natural killer T cells (NKT), group 3 innate lymphoid cells (ILC3) or TCRγδ T cells known to contribute to immediate-early IL-17A responses, were moderately affected. Notably, the Th17 lymphocytes were associated with increased surface expression of CD69, a marker specific of tissue-resident memory (TRM) T lymphocytes in lungs^57^. Finally, depletion of CD4^+^ T lymphocytes was associated with reduced protection of the intranasal LafB vaccine against pneumococcal disease (**Figure 4f**). Thus, an intranasal vaccine composed of LafB antigen and mucosal adjuvant induced protection dependent on lung Th17 lymphocytes with TRM features.

### Healthy human individuals develop LafB-specific immunity

To examine whether LafB might be a suitable vaccine antigen for humans and antigenic in man, we screened plasma from >100 healthy human adults for antigen-specific antibodies. Diphtheria toxoid was used as a positive control. As shown in **Figure 5a**, healthy individuals were all strongly immunoreactive to the diphtheria toxoid. In contrast, LafB-specific antibody responses were rather low using ELISA. However, 10% of individuals demonstrated a stronger antibody response specific for LafB. In addition, using immunoblotting, we found that strong immunoreactivity was associated to LafB detection (**Figure 5b**). Finally, peripheral blood mononuclear cells (PBMC) from healthy donors were isolated and stimulated with recombinant LafB or were incubated with T-cell stimulant phytohemagglutinin (PHA) as a positive control (**Figure 5c**). LafB significantly stimulated IFNγ secretion compared to controls. It should be noted that LafB is highly conserved in pneumococci (**Fig. S3**), and to a lesser extent to other members of the *mitis* groups such as the commensal *S. mitis*, meaning that the presence of LafB antibodies do not strictly indicate previous pneumococcal carriage or infection. Nevertheless, these data indicate that LafB is antigenic in human and a potential universal pneumococcal vaccine antigen that mobilizes lung resident memory Th17 lymphocytes and protects in the context of preexisting viral infections in mice.

**Figure 5.**
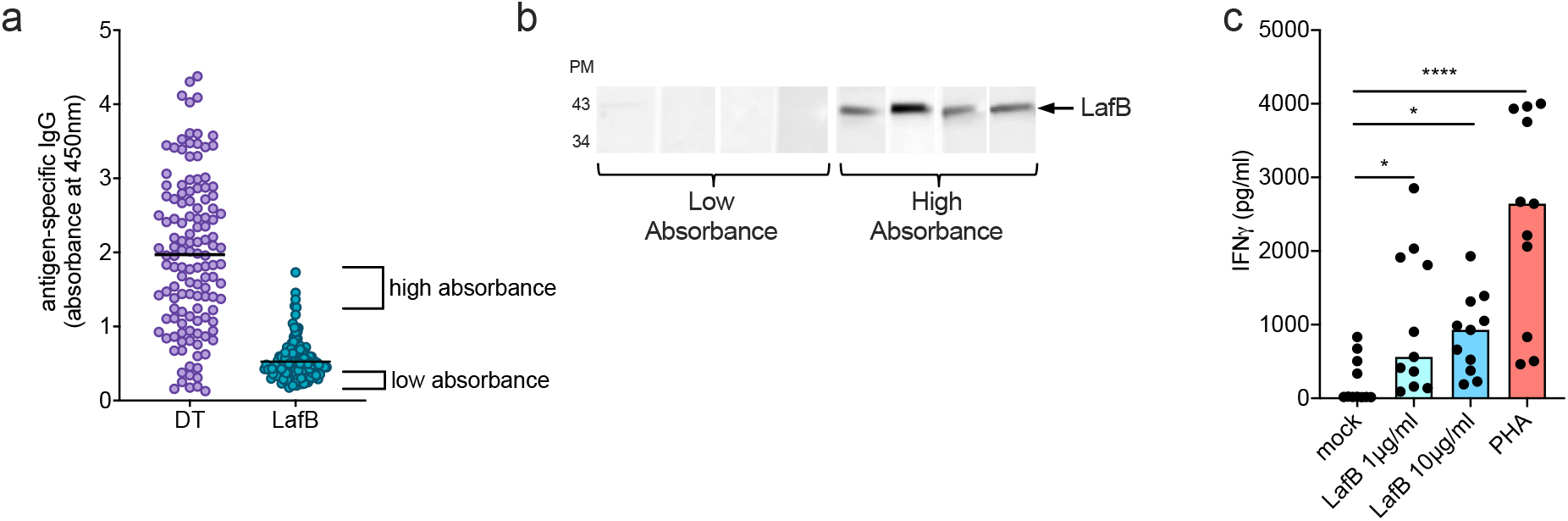
Healthy human individuals have LafB-specific antibody and T cell responses. (**a**) Diphtheria toxoid (DT)- and LafB-specific IgG of plasma from healthy donors (n=127) were determined by ELISA. Plots represent values for individual people as well as median. (**b**) Immunoblot assays of healthy individual plasma specific for LafB. Plasma (n=4/group) with low and high absorbance at 450nm in ELISA were analyzed by immunoblotting. Recombinant LafB was separated by SDS-PAGE and transferred to a membrane before probing with plasma. (**c**) PBMC from healthy donors (n=3) were stimulated 5 days with LafB or PHA or left untreated (mock). The secretion of IFNg was determined by ELISA. Plots represent values for individual values and median. Statistical significance (*p<0.05, *** p<0.001) was assessed by one-way ANOVA Kruskal-Wallis test with Dunn’s correction compared to the medium group.

## Discussion

The principal contribution of this work is the identification of a conserved intracellular membrane-associated pneumococcal antigen as a vaccine candidate effective in protection even following influenza virus infection. The unbiased genetic approach of antigen screening by CRISPRi in the context of superinfection defined that the conserved protein LafB plays an essential role in pneumococcal virulence. LafB is important for proper cell envelope homeostasis, and despite not localized to the bacterial surface and directly exposed to the immune system, the protein triggers vigorous antibody- and T cell-mediated immune responses. This paradigm for antigen selection may open new avenues for discovery of virulence-associated vaccine candidates heretofore overlooked by classical approaches. LafB protection was more effective against pneumococcal challenge when used for intranasal vs. subcutaneous vaccination, but intranasal vaccination did not induce significant circulating or secretory anti-LafB antibodies compared to the subcutaneous route, suggesting that high titer opsonizing antibodies were not pivotal for the protection phenotype. Antigen presentation by the intranasal route may mobilize specific sampling and processing of antigen or unique targeting of antigen-presenting cells coordinating the stimulation of T cell-mediated immunity. In contrast to surface determinants, LafB may only be exposed outside of bacteria upon the production of extracellular vesicles^58^, lysis or autolysis. Indeed, during colonization, pneumococci establish biofilms that consist of a matrix formed from lysed bacterial cells^59^. A subset of healthy humans have measurable LafB-specific IgG levels in their serum, indicating that LafB is also antigenic in human. Deciphering the immune cells and regulatory pathways in host and bacteria involved in the respiratory immune pattern is an important question for future research. In addition, it would be interesting to test whether intranasal vaccination with LafB also protects in a pneumococcal pneumonia model without viral challenge.

Multiple lines of evidence show that Th17 lymphocytes are instrumental for protecting the respiratory mucosa against pneumococcal nasopharyngeal colonization or pneumonia^48–51,60,61^. Moreover, preceding influenza virus infection may blunt subsequent IL-17 production by γδ T cells in response to *S. pneumoniae*^62^. Cross-protection against pneumococcal diseases after recovery from a primary infection is mediated by memory Th17 cells but the antigenic determinants remained to be defined^51^. Interestingly, memory Th17 responses induced by pneumococcal infection can overcome subsequent viral-driven Th17 inhibition and provide cross-protection against different serotypes during coinfection with IAV. Based on these findings, the authors suggested that a vaccine that drives Th17 responses would be potentially able to mitigate disease caused by coinfection^63^. The here discovered, highly conserved LafB may constitute such a prototypic cross-protective antigen. Recent studies highlighted how lung Th17 cells can differentiate into TRM that can persist in tissue, promote long-term robust protection against pathogens^50,57^, and are less prone to alteration or collapse in the context of immunosuppression or immunodysregulation^50^. This unique capacity is noteworthy for protection of high-risk individuals to pneumococcal diseases, such as the elderly, those suffering chronic disease, cancer patients, and transplant recipients, all of whom may be more susceptible to viral infection. Stimulation of mucosal immunity and particularly Th17 lymphocytes and TRM may explain the poorer protective capacity of systemic route of immunization. Similar observations were made for COVID-19 vaccination in which higher antibody levels not correlate with better disease outcome, particularly in older individuals^64^. Our initial experiments using PBMC support LafB as an interesting antigen for human vaccination, but one that will need specific adjuvants to polarize the immunity to Th17 and TRM and target the stimulation and response to key areas of lungs. The use of mucosal adjuvants to potentiate the immune response and, particularly broadly protective lung TRM, is an expanding field of research that will undoubtedly lay the foundation of a new generation of vaccines against respiratory pathogens, including antimicrobial-resistant pathogens^50,65,66^.

## Materials and Methods

Detailed methods are provided in the supporting text.

## Acknowledgments

We thank Dr Mara Baldry and Charlotte Costa for assistance in animal experiments, Dr Frédéric Wallet from Lille Hospital for 15A and 24F clinical isolates, Dr Loic Coutte for *Il17a*^*-/-*^ mice, Mrs Delphine Cayet for LafB-specific ELISA, and Dr Guillaume Lefebvre and Dr Julie Démaret for discussion on pneumonia patients. We thank Dr. Jingwei Xu for his assistance on RoseTTAFold prediction of LafB 3D structure. We also thank UNIL’s Metabolomics and Genomics Technologies Facilty, UAR2014-US41-PLBS, BioImaging Center Lille, Flow Cytometry Core Facility and PLETHA animal facility. This work was supported by an ERC consolidator grant 771534 and the Swiss National Science Foundation (grants 310030_192517, 310030_200792, 51NF40_180541 and IZSEZ0_213879 to J.W.V.; INSERM, Institut Pasteur de Lille, Université de Lille, the Structure Fédérative de Recherche grant PneumoVac to L.V.M.; European Union Horizon 2020 FAIR grant 847786 to J.C.S. and National Nature Science Foundation of China grant 82270012 to X.L.

## Author contributions

Experimentation: XL, LVM, DS, LM, VADS, JD, FPB, MT. Study design and analysis: XL, LVM, LM, VDB, SG, JWV. Writing – original draft: XL, LVM, JCS, JWV. Writing – review & editing: XL, VN, JCS, JWV.

## Data and materials availability

All data are available in the main text or the supplementary materials. CRISPRi-seq data are available at NCBI Sequence Read Archive (SRA) under the following accession number PRJNA895037.

## Supplementary Materials

Supplementary Text

Figs. S1 to S7

Tables S2 to S3

Supplementary References

**Table S1**. Genome-wide fitness values as assessed by CRISPRi-seq of *S. pneumoniae* grown in C+Y vs during superinfection (separate excel file).

**Table S2**. Strains and plasmids used in the study.

**Table S3**. Primers used in the study.

