## Supplementary information for "A conserved antigen induces respiratory Th17-mediated serotype-independent protection against pneumococcal superinfection"

**The PDF file includes:**

Supplementary Text
Figs. S1 to S7
Tables S2 to S3
References

**Other Supplementary Materials for this manuscript include the following:**

Table S1

### Materials and Methods

#### Bacterial strains and culture conditions

All strains, plasmids and primers used are listed in Supplementary Tables 2 and 3 within the supplementary information. *S. pneumoniae* strains were grown in liquid semi-defined C+Y medium, pH=6.8[1] at 37°C from a starting optical density ($OD_{600nm}$) of 0.01 until the appropriate OD. Transformation of *S. pneumoniae* was performed as described previously[2]. Transformants were selected on Columbia agar with 5% sheep blood with antibiotics (100 µg/ml spectinomycin, 250 µg/ml kanamycin, 1 µg/ml tetracycline, 40 µg/ml gentamycin, 0.05 µg/ml erythromycin). *E. coli* strains were grown with LB medium or LB agar with appropriate concentrations of antibiotics (100 µg/ml ampicilline, 100 µg/ml spectinomycin, or 50 µg/ml kanamycin).

#### Phase contrast and fluorescence microscopy

*S. pneumoniae* cells were grown in C+Y medium pH = 6.8 at 37°C to an $OD_{595nm}$ = 0.1 without any inducer and diluted 100 times in fresh C+Y medium supplemented with 100 µM IPTG. After one hour incubation at 37°C, 5% $CO_2$ incubator, collect the cells from 1 ml culture by centrifugation at 8000 g, 2 min. The pellet cells were suspended with 50 µl PBS, and 0.5 µl of the suspension was spotted onto a PBS agarose pad on microscope slides. Visualization of GFP was performed as described previously[3].

#### Split-luciferase HiBiT-tag detection system assay

The assay was performed with the Nano-Glo® HiBiT Extracellular Detection System (Promega). The gene sequence encoding the HiBit tag was inserted to the 5' or 3' end of *lafB* coding sequence by cloning, and the HiBit-tagged *lafB* gene was driven by an IPTG inducible promoter and introduced into the chromosome of *S. pneumoniae* Δ*lafB* mutant. Construction of the mutants is described in the supplementary methods. Note that *lafB* is also known as *cpoA*[4]. The HiBit tag is an 11 amino acid peptide, which can bind to a larger subunit and form a complex with luciferase activity[5]. In this assay, the LgBit subunit can be added into the reaction mixture, but it cannot go through the membrane. So, only when the HiBit tag is exposed to the outside of the cell membrane, luminescence signal can be detected. The *S. pneumoniae* mutants carrying N- or C-terminal Hibit tagged LafB were grown in C+Y acid medium, pH=6.8, at 37°C until

OD600 = 0.3. The culture was then diluted 1:10 into fresh C+Y acid medium with or without 1 mM IPTG for the induction of protein expression. When the OD600 of bacterial culture reached 0.3, the culture was split into two groups: one group was the lysed cell group, in which the bacteria were lysed by addition of 16 µg/ml phage lysin cpl-1[6], and the other group was the intact cell group, without any lysis step performed. The luciferase assay was performed according to the manufacturer's instructions with the lysed and intact cells. 50 µl of bacterial culture was mixed with 50 µl of reaction reagent of the kit in a black 96-wells plate. Bioluminescence was quantified on a Tecan Infinite 200 PRO luminometer at 37°C. Bioluminescence was measured right after the reagent addition. Four replicates for each condition were performed.

**Mouse models**

Mice experiments complied with national, institutional and European regulations and ethical guidelines, and were approved by the Institutional Animal Care and Use Committee (animal facility agreement D59-350009, Institut Pasteur de Lille, protocol reference: APAFIS#16966, 201805311410769_v3). Six to eight weeks old male C57BL/6JRj, *Rorc(γt)-Gfp*$^{TG}$, *Il17a*$^{-/-}$ mice were purchased from Janvier Laboratories (Saint Berthevin, France) or bred and maintained in individually ventilated cages (Innorack® IVC Mouse 3.5) and handled in a vertical laminar flow biosafety cabinet (Class II Biohazard, Tecniplast). All infections were performed in an animal biosafety level 2 facility. For depletion of $CD4^+$ cells, mice received an intraperitoneal injection of 200 µg of GK1.5 monoclonal antibody (rat anti-mouse CD4, Bio X cell) or control isotype on day 34.

**Superinfection assays**

Infections were performed by intranasal (i.n.) route in mice that were previously anesthetized by intraperitoneal (i.p.) injection of 1.25 mg of ketamine and 0.25 mg of xylazine in 200 µL of PBS. Male C57BL/6JRj mice were infected i.n. on day 0 with 50 plaque-forming units (PFU) of the pathogenic murine-adapted H3N2 influenza A virus strain Scotland/20/74 in 30 µL of PBS as previously described[7]. On day 7, mice were infected i.n. with *S. pneumoniae* strains using $5\times10^4$ CFU of the frozen working stocks diluted in 30 µL of PBS or CRISPRi library using $1.5\times10^5$ CFU. For CRISPRi experiments, doxycycline (5mg/kg in 200 µL PBS) or PBS (200 µl) were injected intraperitoneally (i.p.) at 1h before and 9 h after *S. pneumoniae* infection. At 24 h post-pneumococcal infection, mice were euthanized by i.p. injection of 5.47 mg of sodium pentobarbital. Lungs and spleen were collected in 1 ml of PBS, homogenized with a T-25 digital Ultraturrax® (IKA) to evaluate CFU count or extract genomic DNA. In some experiments, mouse survival and weight were recorded every day for 10 days after *S. pneumoniae* infection.

**Bacterial genomic DNA extraction and CRISPRi-seq**

Lung homogenates were mixed with deoxyribonuclease I (10 µg/mL, Sigma-Aldrich) for 10 min at room temperature and filtered through 100 µm meshes. The filtrate was centrifuged at 16,000 *g* and the pellet was used for bacterial gDNA extraction using the NucleoSpin Microbial DNA kit (Macherey-Nagel). For bacterial lysis, 6 cycles of 45 s agitation at a speed of 6 m/s in a FastPrep-24™ homogenizer (MP Biomedicals) were used. DNA was quantified in a NanoDrop spectrophotometer (ThermoFisher) The gDNA was used as template in the one-step PCR to prepare the amplicon libraries for Illumina sequencing as described[2]. Sequencing was performed on an Illumina MiniSeq platform using a 54 dark cycle custom recipe[2]. Raw sgRNA counts were obtained from the fastq files using 2FAST2Q (v.2.4.1) with default settings (PHRED score of minimally 30 and 1 mismatch allowed)[8]. Mouse samples PBS 1 and 6, and DOX 1, 5 and 6 were excluded from downstream analyses, on grounds of low pool diversity even without CRISPRi induction (PBS 6), and bottleneck effects (PBS 1, DOX 1, 5, 6). Samples were excluded if >5% of the strains dropped out of the pool without CRISPRi induction, or if the estimated bottleneck size was smaller than three times the pool diversity. Bottleneck sizes were estimated as before[7], with pneumococcal generation time assumed to be 108 minutes, time of growth 24h, and the total population size at the start $1.5\times10^6$ CFU. Differential fitness analyses were performed using DESeq2 (v.1.34.0) in R (v.4.1.1), with hypothesis testing threshold values of an absolute log2FC>1 and alpha<0.05.

**LafB synthetic lethal screening by CRISPRi-seq**

A *lafB* knockout strain (VL4017) was constructed in the background of P*lac-dcas9* containing strain DCI23[9] as described in the supplementary methods. The pneumococcal sgRNA library (Addgene #170432) was transformed into the resulting strain. Genome-wide fitness quantification via CRISPRi-seq was then performed in both the wild-type and the *lafB* deletion mutant in parallel. Treatment of the libraries was performed as described before[7]. Specifically, the induction treatment with 1 mM IPTG lasted for 21 generations in C+Y medium, and the control group without induction was set up. The bacteria were collected after treatment followed by genomic DNA isolation. CRISPRi-seq was performed as described previously[2]. Raw sgRNA counts were obtained from the fastq files using 2FAST2Q (v.2.4.1) with default settings[8]. Differential fitness analyses were performed using DESeq2 (v.1.34.0) in R (v.4.1.1), with hypothesis testing threshold values of an absolute log2FC>1 and alpha<0.05.

**Mice vaccination**

Mice were vaccinated by the intranasal (i.n.) or subcutaneous (s.c.) route at days 0 and 14. A single vaccine dose per animal contained 20 µg LafB in combination with flagellin $FliC_{\Delta174\text{-}400}$ (2.5 µg) for intranasal vaccination (30µl) or Imject™ Alum (ThermoFisher) for subcutaneous vaccination (100µl). The PPSV vaccine, i.e. Pneumovax® from MSD was used (⅕ human dose/mice). Intranasal vaccination was performed under slight anesthesia by gaseous isoflurane (Axience). Antibody and T cell responses were analyzed on blood, spleen, lung and MdLN at day 28 or at day 35, *i.e.,* 7 days after the influenza virus infection.

**Antigen-specific immune responses**

LafB-specific antibodies in serum were assessed by ELISA. Plates were prepared by absorption of LafB (1 µg/ml in carbonate buffer), overnight at 4 °C on Maxisorp microplates (Nunc), and 1 h blocking at room temperature with 1% dried milk in PBS.

ELISA plates were incubated for 1h at room temperature with serial dilutions of the serum samples. Primary antibody binding was revealed with subsequent incubations with HRP-conjugated goat anti-mouse IgG, IgM or IgA (Southern Biotech) and TMB (BD Bioscience); and measured using a microplate reader at 450/570 nm wavelength. LafB-specific T cell responses were analyzed in spleen, MdLN or lungs. Cells ($1\times10^6$) were incubated for 72h with RPMI 1640 with 10% fetal calf serum, 2mM glutamine, 1mM sodium pyruvate, 10mM HEPES, non-essential amino acids, 100U/100µg Penicillin-Streptomycin and stimulated or not with LafB antigen (50 µg/mL) to measure secretion of IL-13, IL-17A, IL-22, or IFN-$\gamma$ by ELISA.

**Flow cytometry analysis**

Lungs were digested with collagenase IA (Sigma, 1 mg/ml) and DNase I (Sigma, 40µg/ml) during 15 min at 37°C. Cells were separated on Percoll 20% and stained for TCRd-PerCP-eFluor710, CD45-AF700, CD19-, Gr1-APC-Cy7, TCRb-BV421, CD90.2-BV510, NKp46-, CD11b-, CD11c-BV605, CD103-BV711, CD69-PE or CD127-PE-Cy7 (Becton Dickinson or Biolegend). Cells were incubated 4h with Brefeldin A (10µg/ml), PMA (25ng/ml) and ionomycin (500ng/ml) and processed for intracellular staining using the kit Intracellular Fixation & Permeabilization (eBiosciences) and IL-17A-APC or control isotype (REAfinity, Miltenyi Biotec). Data were collected on a BD LSR Fortessa and analyzed with FlowJo software.

**Analysis of plasma and PBMC from healthy individuals**

Plasma (n=127) and whole blood cells (n=3) were collected from healthy donors under strict anonymity (Etablissement Français du Sang "Nord de France", EFS, Lille). Written informed consents were obtained from the donors under EFS contract n°NT/18/2016/200 with respect to Decree n°2007-1220 (articles L1243-4, R1243-61 and following) dated August 10$^{th}$, 2007 of the French Public Health Code (published in the Official Journal of the French Republic of August 14th 2007). The use of human samples was approved by the French Ministry of Education and Research under the agreement DC 2015-2575. LafB-specific plasma were analyzed by ELISA as described above using peroxydase-conjugated goat anti-human IgG antibodies (Sigma-Aldrich). PBMC were purified from blood samples using the SepMate$^{TM}$ (StemCell) as described by the manufacturer. PBMC were cultured (1x10$^6$) for 5 days with RPMI 1640 with 10% fetal calf serum (FCS), 2mM glutamine, 1mM sodium pyruvate, 10mM HEPES, non-essential aminoacids, 100U/100µg Penicillin-Streptomycin and stimulated or not with LafB (1µg/ml or 10µg/ml) or Phytohemagglutinin (PHA, 1µg/ml) to measure secretion of IFN-$\gamma$ by ELISA.

**Rabbit LafB antiserum**

The rabbit antiserum against LafB was produced by Eurogentec with the speedy 28-day program. The protocol uses a non-Freund adjuvant, and the immunization schedule includes 4 injections on days 0, 7, 10 and 19. 100 µg/injection of tag-free LafB protein was used for the immunization. Before the first injection, 1 pre-immune bleed and ELISA were performed to make sure the absence of antibodies against LafB in the naïve rabbit. Then on day 21, 1 medium bleed was performed to test the production of IgG with ELISA, and the final bleed was performed on day 28.

**Western Blot and Immunoblot**

To test the antisera of immunized rabbit or mouse. The *S. pneumoniae* strains were grown in 5 ml of acid C+Y medium to OD600 0.3. The cells were harvested by centrifugation at 8000 g, 5 min. The supernatant was removed and the pellets were resuspended with cold TE buffer (10 ml of 1 M Tris-HCl pH7.5, 2 ml of 0.5 M EDTA pH8, 88 ml of MQ water, to final 100 ml) for 2 times. Resuspend the pellet in 200 µl of TE buffer, and then break down the bacteria by sonication (1s pulse on, 1s pulse off, Amp 40%) until the solution become clear. The total protein concentration was quantified with Biorad protein quantification kit (Biorad, Cat. 5000002), and then all the samples were normalized to the same protein concentration by dilution with TE buffer, which was around 1 mg/ml. 100 µl of cell lysate was mixed with 100 µl of 2×SDS loading buffer, and then boiled at 95°C for 10 min. The samples were centrifuged at 15,000 g for 5 min, and then 10 µl of each sample was loaded into one well of a 12% SDS-PAGE (Bio-Rad, Cat. 4561046). 5 µl of PageRuler Plus Prestained Protein Ladder (Fisher, Cat. 26619) was loaded as marker. The protein samples were transferred onto a PVDF membrane. The PVDF membrane with protein samples was blocked with 5% skim milk (PanReac Applichem, A0830) in PBST (PBS pH7.4 with 0.1% Tween-20) under room temperature for 2 hours. The antisera of rabbit or mouse was diluted 1:500 in PBST and then added onto the membrane for 1 hour incubation at room temperature. The membrane was then washed 3 times, 5 min each time. The secondary antibodies HRP conjugate goat-anti-mouse IgG (Promega, Cat. W4021; 1:2500 dilution in PBST) or HRP conjugate goat-anti-rabbit IgG (Abcam, Cat. AB205718; 1:5000 dilution in PBST) was added onto the membrane incubated with mouse or rabbit antiserum as the first antibody, respectively. The secondary antibody was incubated with the membrane at room temperature for 1 hours, followed by 3 times of washing, 10 min each time. Detection is performed using the SuperSignal West Pico Plus Chemiluminescent Substrate (Thermo scientific, Cat. 34579), and the visualization is performed with the FusionCapt Advance FX7 (Witec AG). To test human plasma, purified LafB antigen (500 ng) was loaded on a 4 to 20% SDS-PAGE, transferred to a nylon membrane, and probed with patient plasma (1:100 dilution) overnight at 4°C. The blot was revealed with a horseradish peroxidase-conjugated goat anti-human IgG secondary antibody (1:10,000 dilution, Sigma-Aldrich) 1h at room temperature and visualized with an enhanced chemiluminescence-based detection kit (West Pico PLUS, Thermo Scientific).

**Doxycycline stock**

Doxycycline hyclate (TCI Europe) stock solutions were prepared in PBS at a concentration of 10 mg/mL, filtered through 0.22 µm sterile membranes, aliquoted and stored at -80°C. Fresh dilutions were prepared from frozen stocks and the doxycycline free base concentration was corrected using the conversion factor 0.8.

**Purification of LafB protein from *E. coli***

The *lafB* gene was cloned with a CPD tag into vector pLIBT7_A and maintained by *E. coli* DH5α. The recombinant vector was transformed into *E. coli* strain BL21 freshly for protein expression. To induce the expression of LafB in *E. coli* BL21, the strain with the recombinant vector was cultured in 500 ml of buffered TB medium to OD600nm ~0.6 at 37°C, 200 rpm. To prepare the buffered TB medium, first make the 10× Phosphate-buffered saline (2.4 g of $KH_2PO_4$ and 12.5 g of $K_2HPO_4$ in 1 L MQ water, autoclave at 121°C, 15 min), and then make the TB medium (24 g of tryptone, 48 g of yeast extract, and 10 ml of glycerol in 900 ml of MQ water, autoclave at 121°C, 15 min), and finally 900 ml of TB medium was mixed with 100 ml of the 10× Phosphate-buffered saline to make 1 L of buffered TB.

When the culture reached OD600nm ~0.6, cool down the culture to 16°C, and then add 0.5 mM IPTG (Isopropyl β-d-1-thiogalactopyranoside) to induce the expression of the recombinant LafB with CPD tag overnight (for ~14 hours). The bacteria were collected by centrifugation at 4°C, 5000 g. The pellets were resuspended with 75 ml of buffer (50 mM Tris-HCl, pH=7.5, 300 mM NaCl, 5% Glycerol, 25 mM Imidazole, 5 mM 2-mercaptoethanol, 1 mM PMSF, 750 Units of nuclease). *E. coli* cells were lysed by sonication. Cell lysates were centrifuged at 18,000 rpm, at 4°C for 30 min. The supernatant was then collected for protein purification with cobalt beads. The protocol for purification of the CPD tagged protein is similar to the protocol published previously[10]. Specifically, the supernatant was directly loaded onto cobalt beads, followed by washing with buffer (20 mM Tris, 100 mM NaCl) to remove the nonspecific bindings. We then used 25 ml of elution buffer (20 mM Tris, 100 mM NaCl with 2 mM inositol hexakisphosphate (InsP6)) to elute the protein. Addition of InsP6 activates the protease activity of CPD and the tag is cleaved off, so the final purified protein is tag free. The elution of LafB protein was further purified with Heparin column, and a gradient washing was made by mixing with buffer A1 (20 mM Tris, 100 mM NaCl) and buffer B1 (20 mM Tris, 1 M NaCl). The purified LafB was checked by SDS-PAGE (Extended Fig. 4).

**Biochemical characterization of the LafB glycosylation activity**

Reactions were done with PBS as principal buffer solution. 30 uL total reaction volumes were set up to contain 10 mM lipid mixture consisting of 1 mM MGlcDAG (1,2-Diacyl-3-alpha-D-glucosyl-sn-glycerol, Avanti) and 9 mM DOPG (1,2-Dioleoyl-sn-glycero-3-phospho-rac-(1-glycerol) sodium salt, Sigma) as well as 1 mM UDP-α-

D-Galactose (Merck Millipore). Volumes of purified LafB to desired concentration were added to start reactions. Samples were incubated at 28 °C for 30 minutes before being quenched by the addition of Methanol to a final concentration of 80%. Samples were flash frozen and kept at -80°C until analysis by mass spectrometry.

**Bacterial colonization**

For quantification of bacteria, lungs and spleen were collected 13 days after infection and homogenized in PBS. Viable counts (colony forming unit [CFU]) were determined by plating serial dilutions onto 5% blood-agar plates.

**Viral RNA quantification**

Total RNA was extracted with the NucleoSpin RNA II kit (Macherey-Nagel, Duren, Germany). For H3N2 RNA detection, 500 ng of total lung RNA were reverse transcribed with Superscript II Reverse Transcriptase (Invitrogen) in the presence of IAV specific primers targeting the segment 7 which encodes the matrix protein 1 (M1) (5' TCTAACCGAGGTCGAAACGTA 3'). The cDNA was amplified by Taqman real-time PCR using Taqman probe FAM-TTTGTGTTCACGCTCACCGTGCC-TAMRA with forward primer: AAGACCAATCCTGTCACCTCTGA and reverse primer: CAAAGCGTCTACGCTGCAGTCC. A plasmid coding the M1 gene was serially diluted to establish a standard curve (Ct values / plasmid copies). Equivalent of 12.5 ng of total lung RNA was thoroughly used to determine the level of M1 RNA in the lung of infected animals by absolute quantification[11].

**Proinflammatory gene expression**

Total RNA was extracted with the NucleoSpin RNA II kit (Macherey-Nagel, Duren, Germany). Total RNA was reverse-transcribed with the High-Capacity cDNA Archive Kit (Applied Biosystems). cDNA was amplified using Takyon™ Low ROX SYBR (Eurogentec). Relative mRNA levels were determined by comparing (a) the cycle thresholds (Ct) for the gene of interest and 2 calibrator genes (ΔCt), *Actb* and *B2m,* and (b) 2–ΔCt values for vaccinated group compared with mock group. The specific primers are CGTCATCCATGGCGAACTG / GCTTCTTTGCAGCTCCTTCGT (*Actb*), TGGTCTTTCTGGTGCTTGTC /GGGTGGCGTGAGTATACTTGAA (*B2m*), CCCTCAACGGAAGAACCAAA / CACATCAGGTACGATCCAGGC (*Cxcl2*), CTCCAGAAGGCCCTCAGACTAC / GGGTCTTCATTGCGGTGG (*Il17a*),

GTTCTCTGGGAAATCGTGGAAA / GTTCTCTGGGAAATCGTGGAAA (*Il6*), and TCAGCAACAGCAAGGCGAAA / CCGCTTCCTGAGGCTGGAT (*Ifng*).

**Strain construction**

*Parent strains for IPTG- or tet- inducible systems.* The prsA-lacI-GmR or prsA-lacI-tetR-GmR fragments were amplified from *S. pneumoniae* strain D-LT-PEP9Ptela[12] with primers OVL1694 and OVL1695. The fragments were then transformed into *S. pneumoniae* and select with Columbia blood agar with 1 µg/ml tetracycline.

*Gene deletions.* Erythromycin resistant marker (eryR) was used as the selection marker for all the deletion mutants in this study. To delete *lafB* gene (SPV_0960), upstream and downstream of *lafB* coding regions were amplified with OVL4184/OVL4185 and OVL4180/4181 oligo pairs, respectively. The DNA sequence containing the coding sequence and ribosome binding site of eryR was amplified with OVL2933/OVL2934. The three amplified upstream, downstream and eryR fragments were then assembled by Golden Gate Assembly using BsaI as the restriction enzyme. The product was then transformed into *S. pneumoniae* and selected with Columbia blood agar with 0.5 µg/ml erythromycin. The procedure for transformation was published by us previously[13]. Deletion of cps locus was performed with the same strategy, and the oligo pairs to amplify the upstream and downstream homologous arms were OVL4610/OVL4611 and OVL4608/OVL4609, respectively.

*Complementary strains.* The complementary strain of *lafB* was made by introducing an ectopic *lafB* driven by its native promoter on ZIP locus of *S. pneumoniae*[14]. The upstream locus of ZIP locus with a spectinomycin resistant marker was amplified from pPEPZ[14] with OVL3252/OVL3253, and the downstream of the ZIP locus was amplified from pPEPZ with OVL3254/OVL3255. The two acquired fragments were then digested with BsmBI. The promoter and coding region of LafB was amplified from the genomic DNA of *S. pneumoniae* D39V with OVL4441 and OVL4442, followed by BsaI digestion. Then the three digested fragments were ligated. The ligation product was then transformed into *S. pneumoniae* and the transformants were selected with Columbia blood agar with 100 µg/ml spectinomycin. As a control for the lafB complementary strain, the empty pPEPZ was transformed into *S. pneumoniae* and the selection was performed in the same way.

*GFP fusion strains.* In this study, a C-terminal and N-terminal GFP fused LafB were constructed both driven by an IPTG-inducible promoter at the ZIP locus. To make the N-terminal version of the fusion, we first amplified 3 fragments. The first was a DNA fragment with the upstream of ZIP, spectinomycin resistant marker specR, Plac promoter, and the msfGFP. The fragment was amplified from pASR108[14] with oligos OVL1841/OVL5845. The second fragment was the LafB coding region without start codon, and was amplified from genomic DNA of *S. pneumoniae* D39V with oligos OVL5846/OVL5847. The third fragment was the downstream of ZIP locus, and was amplified from pASR108 with oligos OVL5851/OVL3255. The three fragments were digested with BsmBI, followed by ligation. The ligation product was transformed into *S. pneumoniae* and 100 µg/ml spectinomycin was used for selection. The C-terminal version of fusion was constructed in a similar way, whereas the oligo pairs OVL1841/OVL5849, OVL5850/OVL5853, OVL5851/OVL3255 were used for amplification of the three fragments.

*IPTG inducible HiBiT tagged lafB mutant.*

Both N- and C-terminal HiBiT-tagged LafB were constructed at the CIL locus[14] under an IPTG-inducible promoter. Firstly, an IPTG-inducible *lafB* was inserted at the CIL locus, and the produced strain was VL4018. To construct VL4018, three fragments were acquired. The first fragment containing upstream of CIL locus and kanamycin resistant marker was amplified from pASR105 with OVL3318/OVL3371, followed by NheI digestion. The second fragment with Plac-lafB was amplified from VL4007 with OVL1754/OVL1225, followed by XhoI and NheI double digestion. The third fragment with downstream of the CIL locus was amplified from pASR105 with oligos OVL925/OVL3321, followed by XhoI digestion. The three digested fragment was then ligated and transformed into *S. pneumonie* VL333 to construct VL4018.

To make the HiBiT-tagged *lafB* on N terminal, two fragments were amplified, BsmI digested and ligated, followed by transformation and selection with 150 µg/ml kanamycin. The first fragment containing upstream of CIL locus, kanamycin resistant marker, and IPTG inducible promoter was amplified from VL4018 with OVL6063/OVL3318. The second fragment containing hibit-tagged *lafB* and downstream of CIL locus was amplified from VL4018 with OVL6064/OVL3321. The C terminal HiBiT-tagged LafB was constructed in a similar way, but OVL3318/6065 and OVL6066/OVL3321 were used to amplify the two fragments from VL4018, respectively.

*Construction of the CRISPRi libraries.* The plasmids with the sgRNA pool were purified with an *E. coli* library (Addgene #170432), and then transformed into different *S. pneumoniae* strains as well described previously[13].

*Construction of luciferase reporter strain (VL2255).* This strain was constructed based on VL2212[15]. The sgRNA targeting *luc* gene was cloned into vector pPEPZ-sgRNAclone (Addgene #141090) as described previously[13], and the produced plasmid was named as pPEPZ-sgRNAluc. The oligos for annealing of the spacer sequence of sgRNAluc were OVL1020/OVL1021. The pPEPZ-sgRNAluc was then transformed into VL2212 and selected with 100 μg/ml spectinomycin on Columbia agar plates.

*Construct the CPD-tagged LafB protein expression plasmid.*
The *lafB* gene was cloned into the plasmid pLIBT7_A_CPDHisOld for the tagging and expression. The *lafB* gene fragment was amplified from genomic DNA of *S. pneumoniae* D39V with two oligos OVL4449/OVL4450, while the backbone of plasmid pLIBT7_A_CPDHisOld was amplified with two oligos "STM121_BsaIins_rev_notag" and "CPDHis_NEW_forward_with_pIDC_overhang". The two fragments were then assembled by Golden Gate Assembly with BsaI as the restriction enzyme. The product was transformed into chemically competent *E. coli* DH5α, and the transformants were selected on LB agar with 100 μg/ml ampicillin. The successfully cloned plasmid "pLIBT7_A_lafB-CPDHisOld" was confirmed by sanger sequencing. For induction of protein expression, the plasmid was transformed into *E. coli* BL21 freshly. The DNA sequence of plasmid "pLIBT7_A_lafB-CPDHisOld" is shown at the end of this document.

*Construction of CRISPRi mutants targeting cozE and divIB.* The sgRNAs targeting *cozE* and *divIB* were cloned into vector pPEPZ-sgRNAclone (Addgene #141090) as described previously [13]. Oligos with the spacer sequences targeting *cozE* and *divIB* were OXL812/OXL813 and OXL814/OXL815, respectively. The pPEPZ-sgRNA vectors with the sgRNAs were then transformed into VL4017 and selected with 100 μg/ml spectinomycin on Columbia agar plates.

**Antisera production in rabbit used for Western blotting**

100 µg/injection of protein was used to raise antibodies in one rabbit, and in total 4 injections were performed on day 0, 7, 10, 19, following the Speedy 28-Day program of Eurogentec. The produced antiserum from the immunized rabbit was shown to be immunogenic against, and specific for pneumococcal LafB by Western blotting.

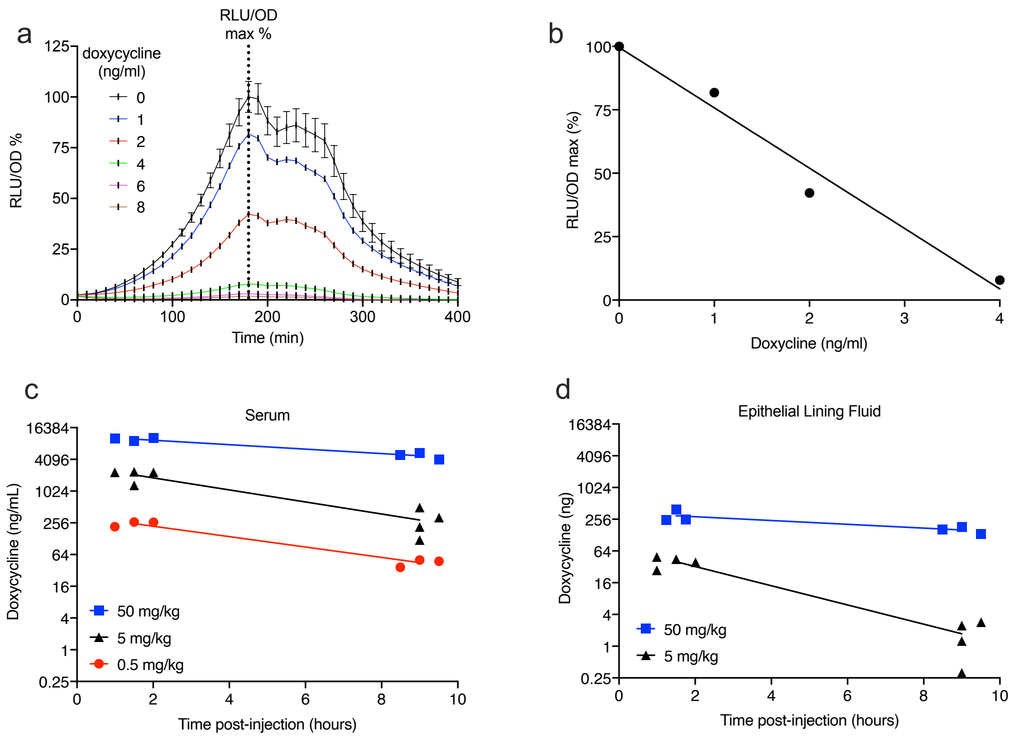
a
RLU/OD
max %
doxycycline
(ng/ml)
0
1
2
4
6
8
RLU/OD %
Time (min)
b
RLU/OD max (%)
Doxycline (ng/ml)
c
Serum
Doxycycline (ng/mL)
50 mg/kg
5 mg/kg
0.5 mg/kg
Time post-injection (hours)
d
Epithelial Lining Fluid
Doxycycline (ng)
50 mg/kg
5 mg/kg
Time post-injection (hours)

**Figure S1. Doxycycline induces *in vitro* and *in vivo* CRISPRi in a dose-dependent manner.** Pneumococcal strain VL2255 was constructed that constitutively expresses firefly luciferase (*luc*), has an sgRNA targeting *luc* and a doxycycline-inducible dCas9. (**a-b**) VL2255 was grown in THYB supplemented with D-luciferin at the indicated concentrations of doxycycline. Luminescence (RLU) and cell density (OD) were measured every 10 minutes in a microplate reader. (**a**) Doxycycline gradually decreases the luminescence of VL2255 strain. Data is presented as a percentage of luminescence compared to the control, *i.e.,* no doxycycline (RLU/OD max %). Mean and SD of two replicates. (**b**) The normalized luminescence at the peak (RLU/OD) has a linear correlation with doxycycline concentration. Data are representative of 1 out of 3 experiments. (**c-d**) C57BL/6 mice (n = 3-4) were injected intraperitoneally with 0.5, 5, or 50 mg/kg of doxycycline, and serum and bronchoalveolar lavages (that represent the epithelial lining fluid or ELF) were sampled at 1.5 h and 9 h. VL2255 was grown in THYB, 10% normal mouse serum, D-luciferin and the various concentrations of serum or lavages from doxycycline-injected animals. (**c**) Concentration of doxycycline in serum. (**d**) Amount of doxycycline in ELF. Data are from one experiment.

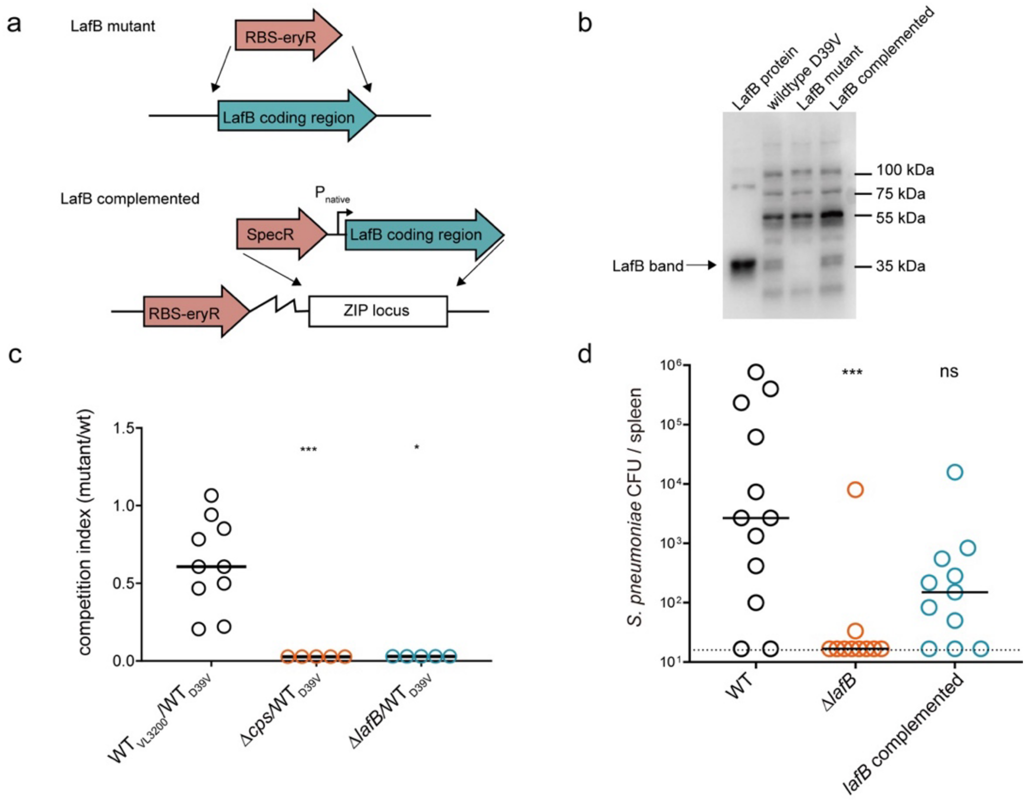
a
LafB mutant
RBS-eryR
LafB coding region
LafB complemented
P native
SpecR
LafB coding region
RBS-eryR
ZIP locus
b
LafB protein
wildtype D39V
LafB mutant
LafB complemented
100 kDa
75 kDa
55 kDa
LafB band
35 kDa
c
competition index (mutant/wt)
***
*
WT VL3200/WT D39V
Δcps/WT D39V
ΔlafB/WT D39V
d
S. pneumoniae CFU / spleen
***
ns
WT
ΔlafB
lafB complemented

**Figure S2. LafB-deficient pneumococci are attenuated for invasive disease.** (**a**) Construction scheme of *lafB* mutant and complemented strain. (**b**) Validation of strains by western blotting using LafB-specific rabbit serum. Purified recombinant tag-free LafB protein (2 ng) was used as a positive control. The cell lysates of wildtype (WT) strain D39V, Δ*lafB* deletion mutant and *lafB* complemented strain as shown in panel (a) were loaded with equal amount of total protein. LafB-specific rabbit serum were generated as described in methods, and HRP-conjugated goat-anti-rabbit IgG was used as secondary antibody. (**c**) Spleen data for competition index of mutant compared to wild type D39V. The same analysis as Figure 1c was performed in spleen. Each dot represents the spleen CFU count at day 8 of a single mouse infected with flu at day 0, and a ratio 1:1 of mutant and WT strain at day 7. Capsule deletion mutant (Δ*cps*) was used as control because capsule is a well-known pneumococcal virulence factor. (**d**) Spleen CFU data for single strain infection (mutant vs complemented vs WT). The same analysis as Figure 1d was performed in spleen. There was a significant difference between the wild-type and *ΔlafB* strain tested by Kruskal-Wallis test. Note that ectopic expression of *lafB* complemented the phenotype of the *lafB* deletion mutant.

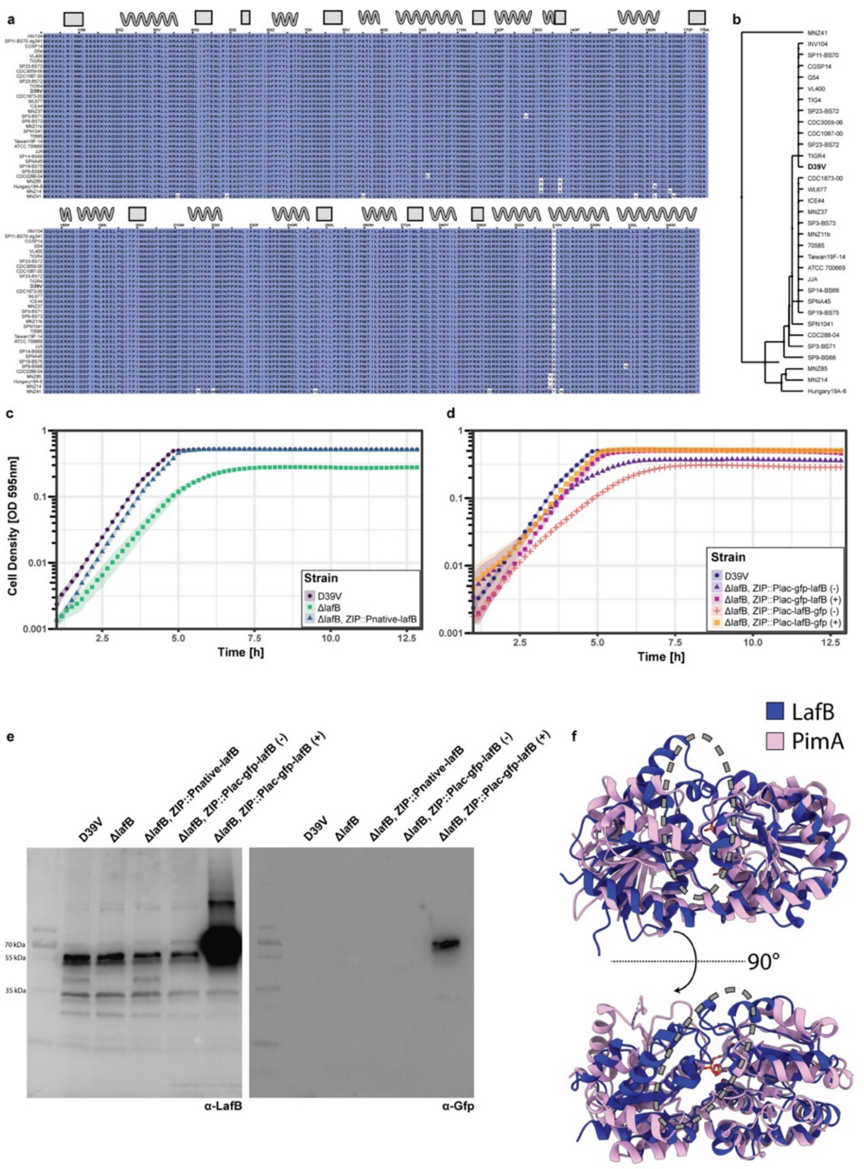
a
b
MNZ41
INV104
SP11-BS70
CGSP14
G54
VL400
TIG4
SP23-BS72
CDC3059-06
CDC1087-00
SP23-BS72
TIGR4
D39V
CDC1873-00
WL677
ICE44
MNZ37
SP3-BS73
MNZ11b
70585
Taiwan19F-14
ATCC 700669
JJA
SP14-BS69
SPNA45
SP19-BS75
SPN1041
CDC288-04
SP3-BS71
SP9-BS68
MNZ85
MNZ14
Hungary19A-6
c
Cell Density [OD 595nm]
Time [h]
Strain
D39V
ΔlafB
ΔlafB, ZIP::Pnative-lafB
d
Strain
D39V
ΔlafB, ZIP::Plac-gfp-lafB (-)
ΔlafB, ZIP::Plac-gfp-lafB (+)
ΔlafB, ZIP::Plac-lafB-gfp (-)
ΔlafB, ZIP::Plac-lafB-gfp (+)
e
D39V
ΔlafB
ΔlafB, ZIP::Pnative-lafB
ΔlafB, ZIP::Plac-gfp-lafB (-)
ΔlafB, ZIP::Plac-gfp-lafB (+)
70 kDa
55 kDa
35 kDa
α-LafB
α-Gfp
f
LafB
PimA
90°

**Figure S3. LafB is a highly conserved membrane-associated pneumococcal GT-B glycosyltransferase.** (**a**) Amino acids sequence alignment of LafB among a selection of relevant *S. pneumoniae* strains spanning a very wide pangenome distribution [16] demonstrating >96% sequence identity. Secondary structure features are indicated above. Background color of each amino acids is based on similarity to the reference sequence of D39V. (**b**) Schematic phylogenetic tree of the LafB proteins in (a). (**c**) Growth curves showing the growth defect of Δ*lafB* strains and the supplementation by expression of LafB from the ZIP locus under the native *lafB* promoter. Growth curves are the result of triplicate measurements with ribbons denoting the confidence interval of the measurement. (**d**) Comparison of growth of strains carrying the GFP-fused LafB proteins with and without supplementation of IPTG in the growth medium demonstrating the supplementation of the LafB phenotype by GFP-tagged LafB proteins. (**e**) Western blots using anti-LafB serum (see Supplementary Information) as well as anti-GFP sera showing the expression of the GFP-LafB protein with and without induction as compared to strains harboring LafB deletion and complementation. GFP-LafB showed the expected size of ~ 65 kDa seemingly lacking any laddering indicative of protein degradation. (**f**) Structural comparison of the predicted structure of LafB and the crystal structure of PimA (PDB: 2GEK). GDP as present in PimA crystal structure was left visible for illustration of substrate binding pocket. Backbone structure of both proteins shows good alignment showing the relation between the protein families with the notable divergence of the active site cleft (indicated by grey dashed oval) as expected due to the different substrate specificities of both proteins.

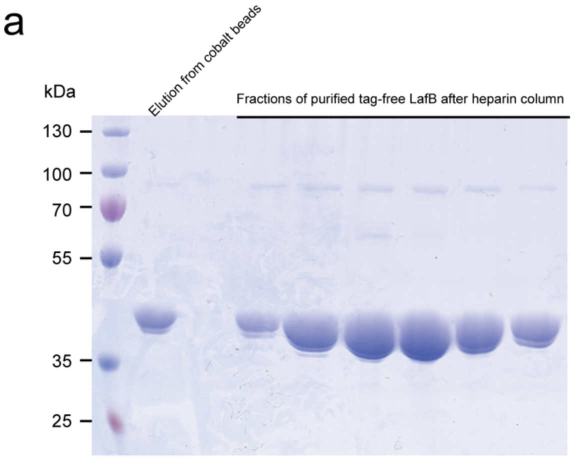

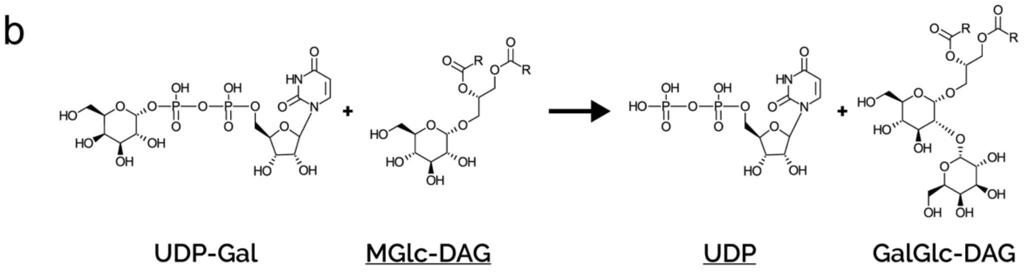

c

| | *Peak Area (Ion counts)* | | |
|---|---|---|---|
| **LafB concentration** | **Uridine** | **UDP-Hexose** | **UDP** |
| 0 μM LafB | 22 | 5086735 | 497 |
| 20 μM LafB | 171 | 3541577 | 43689 |
| 70 μM LafB | 583 | 4506628 | 74045 |

| **LafB concentration** | ***UDP / UDP-Hexose signal ratio*** |
|---|---|
| 0 μM LafB | $9.8 \times 10^{-5}$ |
| 20 μM LafB | 0.012 |
| 70 μM LafB | 0.016 |

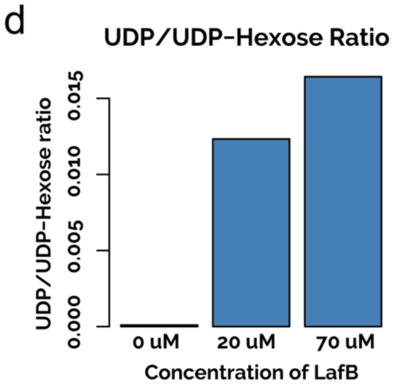

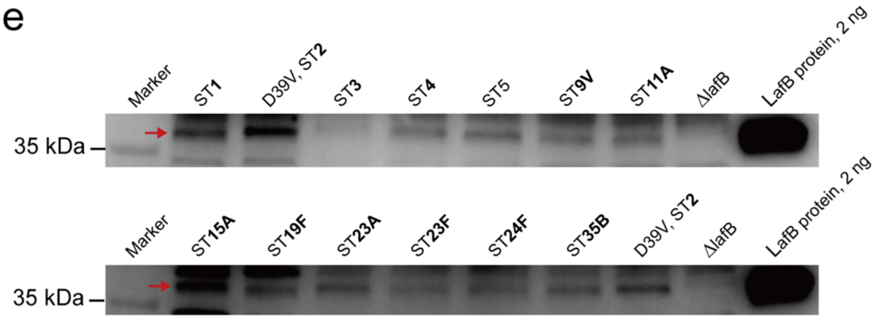

**Figure S4: Purification and biochemical characterization of LafB in *E. coli* and serotype-independent recognition of LafB by serum from LafB-vaccinated mice.** *S. pneumoniae lafB* was cloned in plasmid pLIBT7_A with CPD tag [17], and then the vector was transformed into *E. coli* BL21 for expression. LafB expression was induced by 0.5 mM IPTG in TB medium at 16°C overnight, and then purified with cobalt beads followed by further purification with heparin column. Note that the CPD-His6 tag was cleaved off during purification, see supplementary methods for the details of protein purification. (**a**) Polyacrylamide gel analysis to show the purification of the tag-free LafB. The gel is stained with Coomassie Brilliant Blue. The expected size of tag-free LafB is 40 kDa. (**b**) Schematic of glycosyltransferase reaction of LafB. Underlined compounds were subject to detection via mass spectrometry. UDP-Gal, Uridine-5′-diphosphogalactose; MGlc-DAG, 1,2-diacyl-3-O-(α-D-glucopyranosyl)-sn-glycerol; UDP, Uridine-5′-diphosphate, GalGlc-DAG, 1,2-Diacyl-3-O-[alpha-D-galactopyranosyl-(1->2)-O-alpha-D-glucopyranosyl]-sn-glycerol. (**c**) Mass spectrometry data from LafB reactions. Raw data depicting the total peak area for the indicated chemical species as well as the product to educt signal ratio for the reaction. (**d**) Barplot visualization depicting the product to educt ratio obtained with different concentrations of LafB after incubation. (**e**) Serum from mice vaccinated with LafB and alum (as described in Figure 3a) were used to probe LafB by immunoblotting. Whole bacterial lysates from the following serotypes 1, 2, 3, 4, 5, 9V, 11A, 15A, 19F, 23A, 23F, 24F and 35B were prepared for immunoblotting. As shown, LafB was recognized in all strains by Western blotting, including non-vaccine serotypes 15A and 24F.

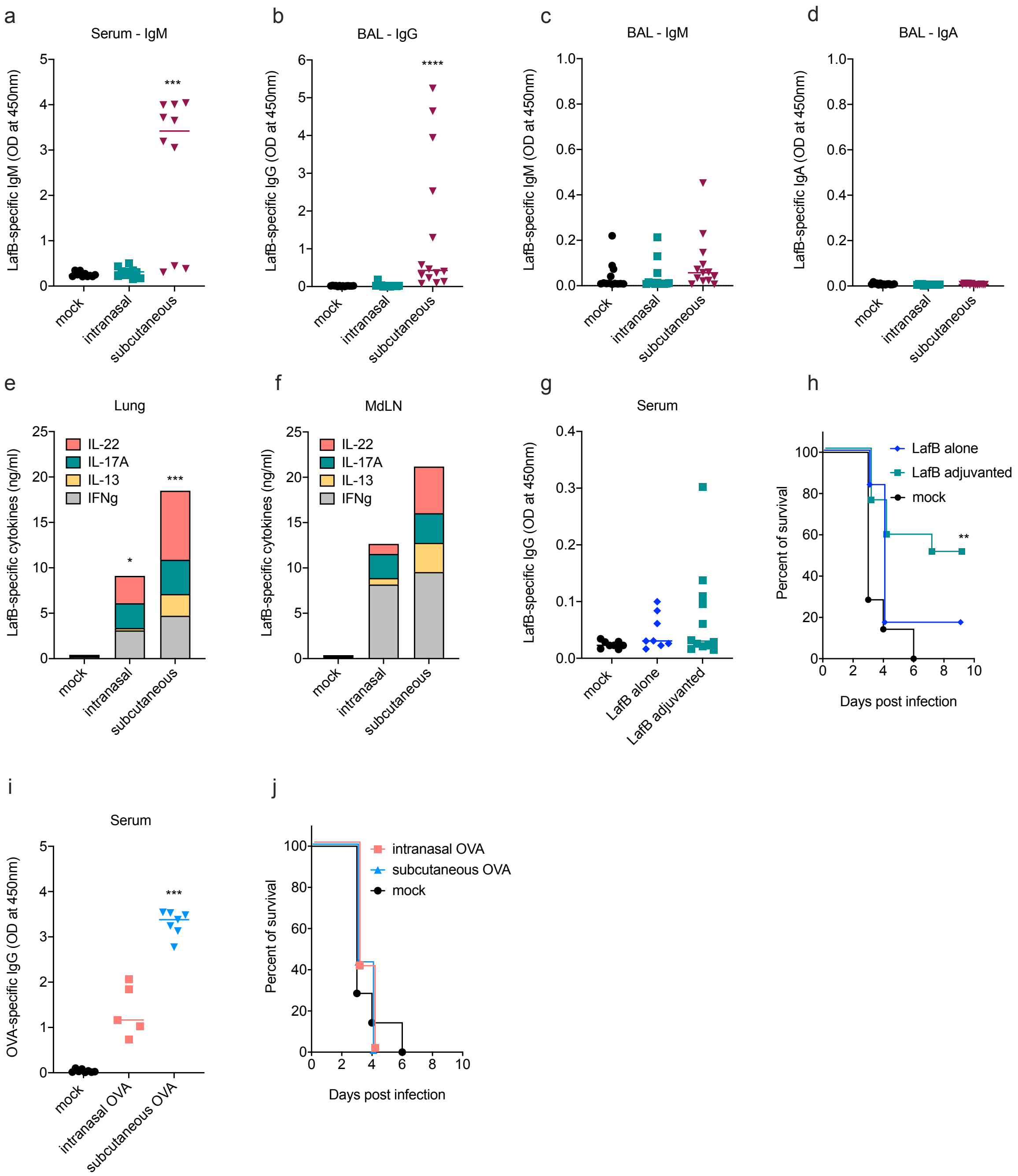
a
Serum - IgM
LafB-specific IgM (OD at 450nm)
mock
intranasal
subcutaneous
***
b
BAL - IgG
LafB-specific IgG (OD at 450nm)
****
c
BAL - IgM
LafB-specific IgM (OD at 450nm)
d
BAL - IgA
LafB-specific IgA (OD at 450nm)
e
Lung
LafB-specific cytokines (ng/ml)
IL-22
IL-17A
IL-13
IFNg
f
MdLN
g
Serum
LafB-specific IgG (OD at 450nm)
LafB alone
LafB adjuvanted
h
LafB alone
LafB adjuvanted
mock
Percent of survival
Days post infection
**
i
Serum
OVA-specific IgG (OD at 450nm)
mock
intranasal OVA
subcutaneous OVA
***
j
intranasal OVA
subcutaneous OVA
mock
Percent of survival
Days post infection

**Figure S5. Protection and immune responses induced by intranasal vaccination depends on the LafB antigen and the mucosal adjuvant flagellin**. (**a-f**) C57BL/6 mice (n=5-15) were immunized at days 0 and 14 with LafB by intranasal (flagellin-adjuvanted) or subcutaneous (alum-adjuvanted) route, or left untreated (mock) and immune responses were analyzed at day 28. (**a-d**) LafB-specific antibody response. (**a**) Serum IgM and (**b-d**) broncho-alveolar lavage (BAL) LafB-specific IgG (**b**), IgM (**c**) or IgA (**d**) were determined by ELISA. (**e-f**) LafB-specific T cell response. Lung (**e**) and mediastinal lymph nodes (MdLN) (**f**) cells were stimulated 72h with LafB and cytokine levels in supernatant were determined by ELISA. (**g-h**) LafB-mediated protection requires mucosal adjuvant. C57BL/6 mice (n=7-11) were immunized at days 0 and 14 by intranasal route with LafB alone, LafB adjuvanted with flagellin or left untreated (mock). (**g**) LafB-specific IgG in serum were determined at day 28 by ELISA. (**h**) Vaccinated mice were infected with influenza A virus at day 28 and were challenged at day 35 with $5\times10^4$ *S. pneumoniae* D39V strain. Protection was assessed by monitoring survival. (**i-j**) Mice were vaccinated with ovalbumin (OVA) as an irrelevant antigen as described in panel a-f. (**i**) OVA-specific IgG response in serum at day 28. (**j**) Protection is specific for LafB antigen. Vaccinated mice were challenged as described in panel h. Protection was assessed by monitoring survival. Plots for antibody represent values for individual mice as well as median. Cytokine data are expressed as median. Statistical significance (*P<0.05, *** p<0.001) was assessed by one-way ANOVA Kruskal-Wallis test with Dunn's correction compared to the mock group. Statistical significance for survival (** p<0.01) was assessed by Mantel-Cox test compared to the mock group.

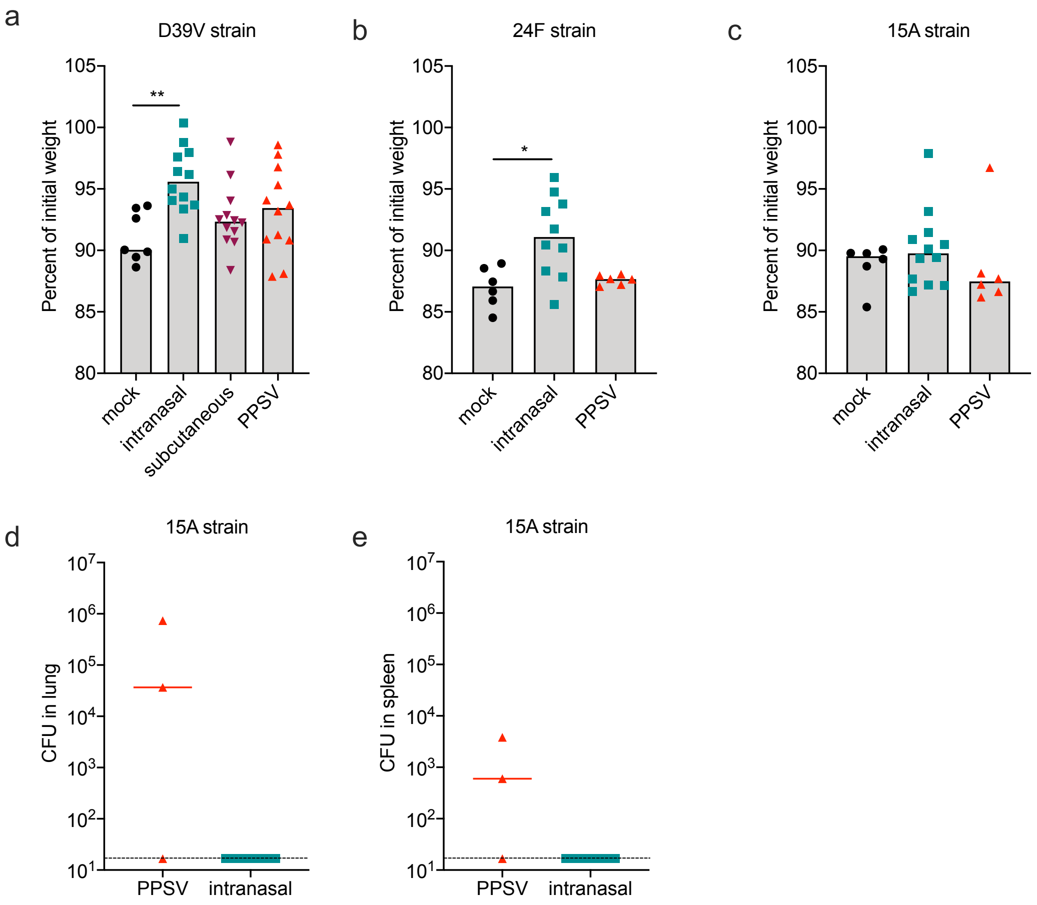
a
D39V strain
Percent of initial weight
105
100
95
90
85
80
**
mock
intranasal
subcutaneous
PPSV
b
24F strain
Percent of initial weight
*
mock
intranasal
PPSV
c
15A strain
Percent of initial weight
mock
intranasal
PPSV
d
15A strain
CFU in lung
$10^7$
$10^6$
$10^5$
$10^4$
$10^3$
$10^2$
$10^1$
PPSV
intranasal
e
15A strain
CFU in spleen
PPSV
intranasal

**Figure S6. Intranasal vaccination with adjuvanted LafB attenuated the pathology caused by *S. pneumoniae* and led to complete bacterial clearance.** C57BL/6 mice (n=6-12) were immunized at days 0 and 14 with adjuvanted LafB intranasal (flagellin-adjuvanted) or subcutaneous (alum-adjuvanted) route, a commercial PPSV vaccine, or untreated (mock). Mice were infected with influenza virus at day 28 and with *S. pneumoniae* at day 35 of the strain D39V of serotype 2 (**a**, $5\times10^4$ CFU), serotype 24F (**b**, $10^3$ CFU), or serotype 15A strain (**c-e**, $5\times10^4$ CFU). (**a-c**) Weight loss at day 36 (24 h post-challenge). Plots represent values for individual mice as well as median. Statistical significance (*P<0.05, ** p<0.01) was assessed by one-way ANOVA Kruskal-Wallis test with Dunn's correction compared to the mock group. (**d-e**) Bacterial load at day 49 (i.e., 14 days post-pneumococcal infection) in lung (**d**) and spleen (**e**) of survivors.

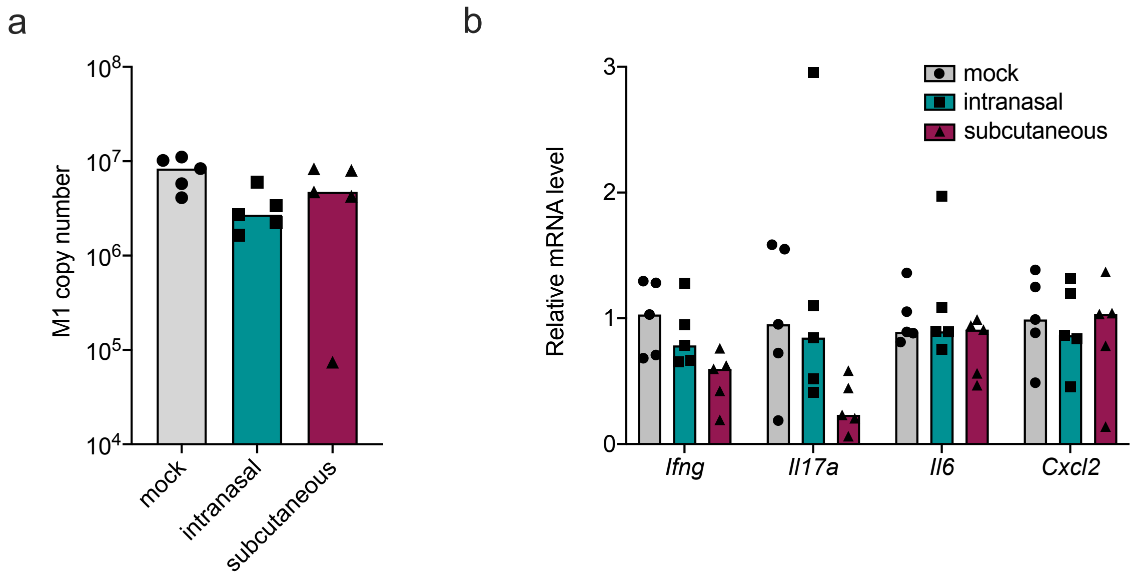
a
M1 copy number
10^8
10^7
10^6
10^5
10^4
mock
intranasal
subcutaneous
b
Relative mRNA level
3
2
1
0
mock
intranasal
subcutaneous
Ifng
Il17a
Il6
Cxcl2

**Figure S7. LafB immunization does not alter the course of lung infection or inflammatory response induced by influenza virus**. C57BL/6 mice (n=5) were immunized or left unvaccinated (mock) at days 0 and 14 with LafB by intranasal (flagellin-adjuvanted) or subcutaneous (alum-adjuvanted) route or left unvaccinated (mock). Mice were infected with H3N2 influenza A virus at day 28 and lungs were sampled seven days later (day 35) for gene expression analysis by qPCR. (**a**) Relative viral RNA level (M1 RNA copies/µg RNA) in lung. (**b**) Expression of pro-inflammatory genes in lung. Plots represent values for individual mice as well as median.

**Supplementary Table 1. Genome-wide fitness values as assessed by CRISPRi-seq of *S. pneumoniae* grown in C+Y vs during superinfection (separate excel file).**

**Supplementary Table 2. Strains and plasmids used in the study**

| **Strains/Plasmids** | **Genotype** | **Reference** |
|---|---|---|
| ***S. pneumoniae*** | | |
| D39V | Serotype 2 strain, wild-type | [18] |
| DCI23 | D39V, Δ*bgaA*::*Plac-*dcas9*sp (tet$^R$); Δ*prs1*::PF6-*lacI* (Gm$^R$ ) | [19] |
| VL4181 | Serotype 15A strain, wild-type, clinical isolate | This study |
| VL4182 | Serotype 24F strain, wild-type, clinical isolate | This study |
| VL1310 | Serotype 1 strain, wild-type, PMEN28, Sweden$^1$-28 Clone, clinical isolate | This study |
| VL1311 | Serotype 3 strain, wild-type, PMEN31, Netherlands$^3$-31 clone, clinical isolate | This study |
| VL2177 | Serotype 4 strain, wild-type, TIGR4 | Sven Hammerschmidt group collection |
| VL1308 | Serotype 5 strain, wild-type, PMEN19, Colombia$^5$-19 Clone, clinical isolate | This study |
| VL1307 | Serotype 9V strain, wild-type, PMEN3, Spain$^{9V}$-3 clone, clinical isolate | This study |
| VL1313 | Serotype 11A strain, wild-type, clinical isolate | This study |
| VL3483 | Serotype 19F strain, wild-type, clinical isolate | This study |
| VL3411 | Serotype 23A strain, wild-type, clinical isolate | This study |
| VL1306 | Serotype 23F strain, wild-type, PMEN1, Spain$^{23F}$-1 clone, clinical isolate | This study |
| VL4304 | Serotype 35B strain, wild-type, clinical isolate | This study |
| VL2212 | D39, Δ*prs*1::PF6-*tetR* (Gm$^R$), Δ*bgaA*:: *Ptet-*dcas9* (tet$^R$) | [15] |
| VL2255 | D39, Δ*prs1*::PF6-*tetR*(Gm$^R$), Δ*bga*::Ptet-*dcas9*(tet$^R$), CIL::P3-*luc* (Kan$^R$), ZIP::P3-sgRNAluc(Spec$^R$) | This study |
| VL3200 | D39V, *hlpA*::*hlpA-mKate-eryR*(ery$^R$) | Veening lab collection |
| VL3508 | D39V, Δ*cps*::*eryR*(ery$^R$) | [15] |
| VL3458 | D39V, Δ*lafB*::*eryR*(ery$^R$) | This study |
| VL3511 | D39V, Δ*lafB*:*eryR*(ery$^R$), ZIP::Plac-empty(Spec$^R$) | This study |
| VL3516 | D39V, Δ*lafB*:*eryR*(ery$^R$), ZIP::Pnative-*lafB*(Spec$^R$) | This study |
| VL333 | D39V, Δ*prs1*::*tetR-lacI* (Gm$^R$) | This study |
| VL4008 | D39V, Δ*prs1*::*tetR-lacI* (Gm$^R$), *lafB*::eryR | This study |
| VL4006 | D39V,Δ*prs1*::*tetR-lacI*(Gm$^R$),ZIP::Plac-msfGFPopt-*lafB*, Δ*lafB*::*eryR*(ery$^R$) | This study |
| VL4007 | D39V, Δ*prs1*::*tetR-lacI*(Gm$^R$), ZIP::Plac-*lafB*-msfGFPopt(Spec$^R$), Δ*lafB*::*eryR*(ery$^R$) | This study |
| VL4017 | D39V, Δ*prs1*::*tetR-lacI*(Gm$^R$), Δ*lafB*::*eryR*(ery$^R$) | This study |

| | | |
|---|---|---|
| VL4018 | D39V, Δ*prs1*::*tetR-lacI*(Gm$^R$), CIL::Plac-*lafB* (Kan$^R$) | This study |
| VL4039 | DCI23, Δ*lafB*::*eryR*(ery$^R$) | This study |
| VL4048 | DCI23, ZIP:sgRNA1-1499 (Spec$^R$) | This study |
| VL4042 | DCI23, Δ*lafB*::*eryR*(ery$^R$), ZIP:sgRNA1-1499 (Spec$^R$) | This study |
| VL4056 | D39V, Δ*prs1*::*tetR-lacI* (GmR), Δ*lafB*::*eryR*, CIL::Plac-Hibit-*lafB* (Kan$^R$) | This study |
| VL4057 | D39V, Δ*prs1*::*tetR-lacI* (Gm$^R$), Δ*lafB*::*eryR*(ery$^R$), CIL::Plac-*lafB*-Hibit (Kan$^R$) | This study |
| SZU486 | D39V, Δ*prs1*::*tetR-lacI*(Gm$^R$), Δ*lafB*::*eryR*(ery$^R$), ZIP-sgRNA*cozE* | This study |
| SZU487 | D39V, Δ*prs1*::*tetR-lacI*(Gm$^R$), Δ*lafB*::*eryR*(ery$^R$), ZIP-sgRNA*divIB* | This study |
| ***E. coli*** | | |
| DH5a | F$^-$*endA1 glnV44 thi-1 recA1 relA1 gyrA96 deoR nupG purB20* φ80d*lacZ*ΔM15 Δ(*lacZYA-argF*)U169, hsdR17(*r$_K^-$m$_K^+$*), λ$^-$ | Veening lab collection |
| BL21 | | Stephan Gruber lab collection |
| **Plasmids** | | |
| sgRNA pools | pPEPZ-sgRNA1-1499 for *S. pneumoniae* D39V (Spec$^R$) | Addgene #170432 |
| pLIBT7_A_CPDHisOld | pUCori-rop-lacI-Plac- CPD10His-f1 ori-AmpR (Amp$^R$) | Stephan Gruber lab collection |
| pLIBT7_A_lafB-CPDHisOld | pUCori-rop-lacI-Plac-lafB-CPD10His-f1 ori-AmpR(Amp$^R$) | This study |
| pPEPZ-sgRNAclone | Addgene # 141090 | [15] |
| pPEPZ-sgRNAluc | with an sgRNA targeting *luc* gene on pPEPZ-sgRNAclone (Spec$^R$) | This study |
| pPEPY (pASR105) | Intergration vector at CIL locus (Kan$^R$) | [14] |
| pASR108 | Intergration vector at ZIP locus with msfGFP (Spec$^R$) | [14] |
| pPEPZ | Intergration vector at ZIP locus (Spec$^R$) | [14] |

Amp$^R$: ampicillin resistant; ery$^R$: erythromycin resistant; Gm$^R$: gentamycin resistant; Kan$^R$: kanamycin resistant; Spec$^R$: spectinomycin resistant; tet$^R$: tetracycline resistant

**Supplementary Table 3. Primers used in this study**

| Primer name | 5'-3' |
|---|---|
| **Primers for gene deletion mutants** | |
| OVL2933_eryR-F-BsaI | GATCGGTCTCGAGGAATTTTCATATGAACAAAAATATAAAATATTCTCAA |
| OVL2934_eryR-R-BsaI | GATCGGTCTCGTTATTTCCTCCCGTTAAATAATAGATAACTATTAAAAAT |
| OVL4180_lafB-dnF-BsaI | gatcggtctcgATAAaaagtgaggtaatctatgcgaatt |
| OVL4181_lafB-dnR | ccagtaggaatgacccg |
| OVL4182_lafB-seqF | gactagatgatggaaaaatgcgc |
| OVL4183_lafB-seqR | tacgcagttctttcaaattttcctg |
| OVL4184_lafB-upF | ggggaaagtctttaacgtaactc |
| OVL4185_lafB-upR-BsaI | gatcggtctcaTCCTtaactactattatatcattttctttg |
| OVL4608_cps-dn-F-BsaI | GATCGGTCTCAATAAgttgtttgaaaaataattttcaaaaattctg |
| OVL4609_cps-dnR | attatctgataaatcccagtcttcgc |
| OVL4610_cps-up-R-BsaI | gatcggtctcTTCCTatacattgaacatcttacgattatatcactttttta |
| OVL4611_cps-upF | ctccctcgtattgtctcaatct |
| **Primers for construction of complementary strains** | |
| OVL3252_ZIP-upF | GCCAATAAATTGCTTCCTTGTTTTG |
| OVL3253_ZIPup-R-BsmBI | ccttcgtctcgACTAGTGAATTCTATAAACGCAGAAAG |
| OVL3254_ZIPdn-F-BsmBI | ccttcgtctcgAGGAAAAATAATGCCGGATCCCT |
| OVL3255_ZIPdn-R | ATGACACGGATTTTAAGAATAATTCTTTCT |
| OVL4441_lafBcom-F-BsaI | gatcggtctcCTAGTacatataaaagcatgtgagagactgttgg |
| OVL4442_lafBcom-R-BsaI | gatcggtctcTTCCTagattacctcactttttactttctccc |
| **GFP fusions** | |
| OVL1841_pPEPZ_F | ACAAAAGTGTGCTATTCTTTTTATGAGAG |
| OVL5845_msfGFP-R-BsmBI | tcagcgtctccTTTGTATAGTTCGTCCATGCC |
| OVL5846_linker-lafB-F | GATCCGTCTCACAAAggatccggatctggtggagaagctgcagctaaaggatcagagaaaaagaaattacgcatcaat |
| OVL5847_lafB-R-BsmBI | gatccgtctctATTActttctccctaaagcggc |
| OVL3255_ZIPdn-R | ATGACACGGATTTTAAGAATAATTCTTTCT |
| OVL5849_pASR108-up-R-BsmBI | gatccgtctcaccatATTTGCCTCCTTAAAGATCTTAATTG |
| OVL5850_lafB-RBS-F-BsmBI | gatccgtctcgatggagaaaaagaaattacgcatca |

| | |
|---|---|
| OVL5851_msfGFP-linker-F-BsmBI | gatccgtctcaggatccggatctggtggagaagctgcagctaaaggatcaTCAAAAGGCGAAGAACTATTCACA |
| OVL5853_msfGFP-eryR-R-BsmBI | gatccgtctcaTCCTTTATTTGTATAGTTCGTCCATGC |
| OVL6159_cozEa-F-Nter-BsmBI | gatccgtctcgatcatttcgtagaaataaattattttttggacc |
| OVL6160_cozEa-R-Nter-BsmBI | gatccgtctcaCGAGTTActtagctaattctctttctcgttc |
| OVL6164_cozEa-F-Cter-BsmBI | gatccgtctcgagttatgtttcgtagaaataaattattttttggac |
| OVL6165_cozEa-R-Cter-BsmBI | GATCCGTCTCACCAGcttagctaattctctttctcgttct |
| **HiBiT tag fusions** | |
| OVL6063_cil-up-Hibit-R-bsmbI | gatccgtctctACCAGAAATTTTTTAAACAAACGCCAACCAGAAACcataactactCCTCCTGATCTTAATTGTG |
| OVL6064_lafB-Hibit-F-bsmbI | GATCCGTCTCACTGGTGGTGGTGGTTCTGGTGGTGGTGGTTCTgagaaaaagaaattacgcatcaatatgt |
| OVL6065_lafB-Hibit-R-bsmBI | GATCCGTCTCAACCAGAAACAGAACCACCACCACCAGAACCACCACCACCctttctccctaaagcggct |
| OVL6066_hibit-cil-dn-F-bsmBI | gatcCGTCTCATGGTTGGCGTTTGTTTAAAAAATTTCTTAATAACTCGAGAAAGTGTAAGCAATTCTG |
| OVL3318_pASR105_UpF | AAACCTTACTAAAGTATATAATTTAGGC |
| OVL3371_(pPEPY)-SPV_564*mNeon_R | AATCCGCGTCTCCcatTAATTTTCCTCCTTATTTATTTAGATCTCATGAATTCAATTGTG |
| OVL1754_LK354 | TGGTCTCCGATCGTGTACTG |
| OVL1225_ | GATTGCCCTCTTGTTCAG |
| OVL925_pPEPY-Linear-R | GGATCCCTCCAGTAACTCGAGAA |
| OVL3321_pASR105_DoR | ATAAAAACATTCATCATAACCCCC |
| **CRISPRi targeting *cozE* and *divIB*** | |
| OXL812_sgRNA-Sp-cozE-F | tataGGTTAAGAGTAAAATTTCTG |
| OXL813_sgRNA-Sp-cozE-R | aaaccagaaattttactcttaacc |
| OXL814_sgRNA-Sp-divIB-F | tataTAACTCTTTCAATTCTTCGA |
| OXL815_sgRNA-Sp-divIB-R | aaactcgaagaattgaaagagtta |
| **Insertion of *tetR* and *lacI*** | |
| OVL1694_prsA lacI-tetR-GmR FWD | AGGACACACCTGCAGTGCCTTATTATTATTGTCCACTTTCCAAAC |

| | |
|---|---|
| OVL1695_prsA lacI-tetR-GmR REV | AGGACACACCTGCAGTGTATGCTCGAAGATTTCAGCTTGACATT |
| **Construction of LafB protein expression plasmid** | |
| STM121 BsaIins rev notag | acgtatggtctccatggttatatctccttcttaaagttaaacaaaattatttctagaggg |
| CPDHis NEW forward with pIDC overhang | CTCCGGAATATTAGGTCTCAggatctCTCGCGGGCGGTAAAATACTCC |
| OVL4449_lafB-TMA-F-BsaI | ctccggaatattaggtctcaCCATggagaaaaagaaattacgcatcaatat |
| OVL4450_lafB-TMA-R-BsaI | GGCTCAAGCAGTGGGTCTCCATCCctttctccctaaagcggc |
| **Construction of the luciferase reporter strain** | |
| OVL1020_GG-sgRNAluc-F | AAACGGCGCCATTCTATCCTCTAG |
| OVL1021_GG-sgRNAluc-R | TATACTAGAGGATAGAATGGCGCC |

**DNA Sequence of plasmid pLIBT7_A_lafB-CPDHisOld**

Note : start codon (ATG) and stop codon (TAA) of the *lafB*-CPD coding sequence was labelled.

tattttctccttacgcatctgtgcggtatttcacaccgcatatatggtgcactctcagtacaatctgctctgatgccgcatagttaagccagta
tacactccgctatcgctacgtgactgggtcatggctgcgccccgacacccgccaacacccgctgacgcgccctgacgggcttgtctg
ctcccggcatccgcttacagacaagctgtgaccgtctccgggagctgcatgtgtcagaggttttcaccgtcatcaccgaaacgcgcga
ggcagctgcggtaaagctcatcagcgtggtcgtgaagcgattcacagatgtctgcctgttcatccgcgtccagctcgttgagtttctcca
gaagcgttaatgtctggcttctgataaagcgggccatgttaagggcggttttttcctgtttggtcactgatgcctccgtgtaagggggattt
ctgttcatgggggtaatgataccgatgaaacgagagaggatgctcacgatacgggttactgatgatgaacatgcccggttactggaac
gttgtgagggtaaacaactggcggtatggatgcggcgggaccagagaaaaatcactcagggtcaatgccagcgcttcgttaatacag
atgtaggtgttccacagggtagccagcagcatcctgcgatgcagatccggaacataatggtgcagggcgctgacttccgcgtttccag
actttacgaaacacggaaaccgaagaccattcatgttgttgctcaggtcgcagacgttttgcagcagcagtcgcttcacgttcgctcgcg
tatcggtgattcattctgctaaccagtaaggcaaccccgccagcctagccgggtcctcaacgacaggagcacgatcatgcgcacccgt
ggggccgccatgccggcgataatggcctgcttctcgccgaaacgtttggtggcgggaccagtgacgaaggcttgagcgagggcgt
gcaagattccgaataccgcaagcgacaggccgatcatcgtcgcgctccagcgaaagcggtcctcgccgaaaatgacccagagcgc
tgccggcacctgtcctacgagttgcatgataaagaagacagtcataagtgcggcgacgatagtcatgccccgcgcccaccggaagg
agctgactgggttgaaggctctcaagggcatcggtcgagatcccggtgcctaatgagtgagctaacttacattaattgcgttgcgctcac
tgcccgctttccagtcgggaaacctgtcgtgccagctgcattaatgaatcggccaacgcgcggggagaggcggtttgcgtattgggc
gccagggtggtttttcttttcaccagtgagacgggcaacagctgattgcccttcaccgcctggccctgagagagttgcagcaagcggtc
cacgctggtttgccccagcaggcgaaaatcctgtttgatggtggttaacggcgggatataacatgagctgtcttcggtatcgtcgtatcc

cactaccgagatatccgcaccaacgcgcagcccggactcggtaatggcgcgcattgcgcccagcgccatctgatcgttggcaacca
gcatcgcagtgggaacgatgccctcattcagcatttgcatggtttgttgaaaaccggacatggcactccagtcgccttcccgttccgcta
tcggctgaatttgattgcgagtgagatatttatgccagccagccagacgcagacgcgccgagacagaacttaatgggcccgctaaca
gcgcgatttgctggtgacccaatgcgaccagatgctccacgcccagtcgcgtaccgtcttcatgggagaaaataatactgttgatgggt
gtctggtcagagacatcaagaaataacgccggaacattagtgcaggcagcttccacagcaatggcatcctggtcatccagcggatagt
taatgatcagcccactgacgcgttgcgcgagaagattgtgcaccgccgctttacaggcttcgacgccgcttcgttctaccatcgacacc
accacgctggcacccagttgatcggcgcgagatttaatcgccgcgacaatttgcgacggcgcgtgcagggccagactggaggtggc
aacgccaatcagcaacgactgtttgcccgccagttgttgtgccacgcggttgggaatgtaattcagctccgccatcgccgcttccactttt
tcccgcgttttcgcagaaacgtggctggcctggttcaccacgcgggaaacggtctgataagagacaccggcatactctgcgacatcgt
ataacgttactggtttcacattcaccaccctgaattgactctcttccgggcgctatcatgccataccgcgaaaggttttgcgccattcgatg
gtgtccgggatctcgacgctctcccttatgcgactcctgcattaggaagcagcccagtagtaggttgaggccgttgagcaccgccgcc
gcaaggaatggtgcatgcaaggagatggcgcccaacagtcccccggccacggggcctgccaccatacccacgccgaaacaagcg
ctcatgagcccgaagctcttcccgataatacgactcactataggggaattgtgagcggataacaattcccctctagaaataattttgtttaa
ctttaagaaggagatataaCCATGgagaaaaagaaattacgcatcaatatgttgagttcaagtgagaaagtagcaggacagggag
tttcaggtgcttaccgtgaattagttcgtcttcttcaccgtgctgccaaggaccaattgattgttacagaaaatcttccaatcgaggcagatg
tgactcactttcatacgattgattttccctattatttatcaaccttccaaaagaaacgctcagggagaaagattggctatgtgcatttcttgcc
agctacacttgagggaagtttgaaaattccatttttcttaaagggaattgtgaaacgctatgtattttctttttacaaccggatggagcacttg
gttgtggtcaatcctatgtttattgaggatttggtagcagctggtattccacgtgaaaaagtgacctatattcctaactttgtcaacaaggaa
aaatggcatcctctaccacaagaagaggtagttagactgcgcacagatcttggtcttagtgacaatcagtttatcgtagtaggtgctggg
caagttcagaaacgtaaagggattgatgactttatccgtctggctgaggaattgcctcagattacctttatctgggctggtggcttctctttt
ggtggtatgacagatggttatgaacactataagaaaattatggaaaatccccctaaaaatttgatttttccaggcattgtatcgccagagc
ggatgcgcgaattgtatgctctagcggatcttttcttgttgcctagttacaatgagctctttcctatgactattttagaagctgcgagttgtga
ggctcctattatgttgcgtgatttagatctctataaggtgattttggagggaaattatcgggcgacagcgggtagagaagagatgaaaga
ggctattttggaatatcaagcaaatcctgctgtcttaaaagatctcaaagaaaaggctaagaatatttccagagagtattctgaagagcat
ctgttacaaatctggttggacttttatgagaaacaagccgctttagggagaaagggatctgatgccctcgcgggcggtaaaatactccat
aatcaaaatgttaatagctggggcccgattacggttacaccaacgacagatggtggtgaaacccgcttcgacggtcaaatcatcgttca
aatggaaaacgacccggtagtagcaaaagcggcagccaatttagcaggtaaacatgctgaaagcagtgtggtggtgcagctcgattc
agacggcaactatcgcgtggtgtatggcgatccgtcaaaactggatggaaagctacgttggcagttggtggggcatggtcgcgacca
ctcagaaactaacaatactcgcttaagtggttacagtgccgatgagttggccgtgaaattggccaagttccaacagtcgtttaatcaagc
cgaaaacatcaacaacaaaccggatcacatcagtattgttggttgttctttggtgagtgacgacaagcaaaaaggctttggtcatcagttt
attaacgcgatggatgcgaatggtcttcgtgtcgatgtctctgttcgtagttctgaactggccgtagacgaggcgggacgtaagcatacc
aaggacgcgaatggcgattgggttcaaaaggcagaaaacaacaaagtttcgctaagctgggacgcgcaaggccttgagcaccacca
ccaccaccaccaccaccaccatTAAgctagcataaccccttggggcctctaaacgggtcttgaggggttttttggggaccggaaga
gcggcgcattaagcgcggcgggtgtggtggttacgcgcagcgtgaccgctacacttgccagcgccctagcgcccgctcctttcgcttt

cttcccttcctttctcgccacgttcgccggctttccccgtcaagctctaaatcgggggctccctttagggttccgatttagtgctttacggca
cctcgaccccaaaaaacttgattagggtgatggttcacgtagtgggccatcgccctgatagacggtttttcgccctttgacgttggagtcc
acgttctttaatagtggactcttgttccaaactggaacaacactcaaccctatctcggtctattcttttgatttataagggattttgccgatttcg
gcctattggttaaaaaatgagctgatttaacaaaaatttaacgcgaattttaacaaaatattaacgtttacaatttcaggtggcacttttcggg
gaaatgtgcgcggaacccctatttgtttatttttctaaatacattcaaatatgtatccgctcatgagacaataaccctgataaatgcttcaata
atattgaaaaaggaagagtatgagtattcaacatttccgtgtcgcccttattcccttttttgcggcattttgccttcctgtttttgctcacccaga
aacgctggtgaaagtaaaagatgctgaagatcagttgggtgcacgagtgggttacatcgaactggatctcaacagcggtaagatcctt
gagagttttcgccccgaagaacgttttccaatgatgagcacttttaaagttctgctatgtggcgcggtattatcccgtattgacgccgggc
aagagcaactcggtcgccgcatacactattctcagaatgacttggttgagtactcaccagtcacagaaaagcatcttacggatggcatg
acagtaagagaattatgcagtgctgccataaccatgagtgataacactgcggccaacttacttctgacaacgatcggaggaccgaagg
agctaaccgcttttttgcacaacatgggggatcatgtaactcgccttgatcgttgggaaccggagctgaatgaagccataccaaacgac
gagcgtgacaccacgatgcctgcagcaatggcaacaacgttgcgcaaactattaactggcgaactacttactctagcttcccggcaac
aattaatagactggatggaggcggataaagttgcaggaccacttctgcgctcggcccttccggctggctggtttattgctgataaatctg
gagccggtgagcgtgggtcccgcggtatcattgcagcactggggccagatggtaagccctcccgtatcgtagttatctacacgacgg
ggagtcaggcaactatggatgaacgaaatagacagatcgctgagataggtgcctcactgattaagcattggtaactgtcagaccaagtt
tactcatatatactttagattgatttaaaacttcatttttaatttaaaaggatctaggtgaagatcctttttgataatctcatgaccaaaatcccttа
acgtgagttttcgttccactgagcgtcagaccccgtagaaaagatcaaaggatcttcttgagatcctttttttctgcgcgtaatctgctgctt
gcaaacaaaaaaaccaccgctaccagcggtggtttgtttgccggatcaagagctaccaactctttttccgaaggtaactggcttcagca
gagcgcagataccaaatactgtccttctagtgtagccgtagttaggccaccacttcaagaactctgtagcaccgcctacatacctcgctc
tgctaatcctgttaccagtggctgctgccagtggcgataagtcgtgtcttaccgggttggactcaagacgatagttaccggataaggcg
cagcggtcgggctgaacggggggttcgtgcacacagcccagcttggagcgaacgacctacaccgaactgagatacctacagcgtg
agctatgagaaagcgccacgcttcccgaagggagaaaggcggacaggtatccggtaagcggcagggtcggaacaggagagcgc
acgagggagcttccagggggaaacgcctggtatctttatagtcctgtcgggtttcgccacctctgacttgagcgtcgatttttgtgatgct
cgtcaggggggcggagcctatggaaaaacgccagcaacgcggcctttttacggttcctggccttttgctggccttttgctcacatgttctt
tcctgcgttatcccctgattctgtggataaccgtattaccgcctttgagtgagctgataccgctcgccgcagccgaacgaccgagcgca
gcgagtcagtgagcgaggaagcggatgagcgcctgatgcgg
